# Identification of Potential inhibitors for Hematopoietic Prostaglandin D2 Synthase: Computational Modeling and Molecular Dynamics Simulations

**DOI:** 10.1101/2021.08.19.456954

**Authors:** Satyajit Beura, Chetti Prabhakar

## Abstract

To design a new therapeutic agent for Hematopoietic Prostaglandin D_2_ synthase (hPGDS), a set of 60 molecules with different molecular scaffolds were (range of pIC_50_ values are from 8.301 to 3.932) considered to create a pharmacophore model. Further, identification of potential hPGDS inhibitors were carried out by using virtual screening with different databases (from 15,74,182 molecules). The Molecular screening was performed using different sequential methods right from Pharmacophore based virtual screening, molecular docking, MM-GBSAstudies, ADME property analysis and molecular dynamics simulations using Maestro11.9 software. Based on the best pharmacophore model (ADRR_1), the resultant set of 18,492 molecules were screened. The preliminarily screened molecules were subjected to molecular docking (PDB_ID: 2CVD) methods. A set of 27 molecules was screened from the resultant molecular docking outcomes (360 molecules) based on binding free energy (ΔG_bind_) and Lipinski’s rule of five. Out of 27 molecules, 4 were selected visual data analysis and further subjected to molecular dynamics (MD) simulation study. Outcomes of the present study conclude with three new proposed molecules (**SP1, SP2** and **SP10)** which show a good range of interaction with human hPGDS enzyme in comparison to the marketed compounds i.e., **HQL-79**, **TFC-007**, **HPGDS inhibitor I** and **TAS-204**.

## 1. Introduction

Prostaglandin D2 (PGD2) is a pro-inflammatory lipid mediator downstream of the cyclooxygenase (COX) pathway [1–3]. Arachidonic acid-derived lipid mediators like leukotrienes, lipoxins, thromboxane A_2_, PGD2, Prostaglandin E2 (PGE2) play a central role in inflammation. Out of different lipid mediators, PGD2 is specifically responsible for allergy development and progression. PGD2 shows its function by activating two-protein-coupled receptors i.e., DP1 (d-type prostanoid receptor 1) and DP2 (d-type prostanoid receptor 2), the latter also being referred to as chemo-attractant receptor homologous-molecule expressed in Th2 cells (CRTH2) [4]. DP1-mediated responses include inhibition of platelet aggregation, bronchodilatation and vasorelaxation [5], and also DP1 antagonists have been found to ameliorate rhinitis, conjunctivitis and pulmonary inflammation in animal models [6–8]. DP2/CRTH2 receptor-mediated response including initiation and potentiation of immune cell migration, respiratory burst, type 2 cytokine productions and histamine release [3]. PGD2 is a potent target for inflammation; its influence strongly depends on whether it acts in the early or late phase of inflammation. On the one hand, in the early phase of inflammation, acute inflammation, i.e., dermatitis [9] and colitis [10], lipopolysaccharide-induced pulmonary inflammation [11] as well as in anaphylactic shock [12], PGD2 seems to have protective effects. On the other hand, in late-phase skin inflammation [9], chronicand allergic inflammation [13–14], PGD2/CRTH2/DP2 activation exacerbates leukocyte migration, activation and survival, while DP1 activation has been linked to increased mucus production and airway hyper reactivity [15]. PGD2 is synthesized by two different enzymes like hematopoietic PGD synthase (hPGDS) and lipocalin-type PGD synthase (LPGDS) [16]. Majorly 90% of PGD2 is synthesized by the enzyme hPGDS. Hematopoietic Prostaglandin D_2_ signaling as a therapeutic target for allergic diseases like allergic asthma, rhinitis [17], atopic dermatitis [18], food allergy, gastrointestinal allergic disorder [19], and anaphylaxis [20].

In the past few decades several research articles and patent works have been introduced for hPGDS inhibitors as a therapeutic option in allergic inflammation, asthma, chronic obstructive pulmonary disease and other inflammatory diseases [21–23]. Some commercially available compounds like HQL-79 [24–26], TFC-007 [27], HPGDS inhibitor I [28], TAS-204 [29], ZL-2102 [30], TAS-205 [31], KMN-698 [32] were used as hPGDS inhibitors. For the treatment of Chronic obstructive pulmonary disease (COPD), asthma and idiopathic pulmonary fibrosis, Phase I clinical trial of hPGDS inhibitor ZL-2102 was initiated in the year of 2015 [30]. Another Phase I clinical trial of hPGDS inhibitor is TAS-205, in which 23 boys with Duchenne’s muscular dystrophy were examined in 2018 [31].

In this study, a set of 60 molecules were considered to create a pharmacophore model that can efficiently explain the essential features required for the inhibition of hPGDS and to generate a model that alleviates in distinguishing molecules that have good efficiency. These different molecular scaffolds inclusive of indole **(A1-8)**, pyridine **(B9-17)** [33], benzaldehyde **(C18-22)** [34], thiophene **(D23-29)** [35], benzimidazole (TAS-204 derivatives) **(E30-41)** and pyrimidine (TFC-007 derivatives) **(F42-60)** [36] based molecules have been taken as a premise against hPGDS in this study (Scheme 1, detailed structures are shown in supporting information Table S1). Ligand-based pharmacophore generation, pharmacophore-based virtual screening, molecular screening, ADME property analysis and molecular dynamics simulations have been employed to study and identify the new candidate having better interaction and binding affinity with the human hPGDS enzyme from the various available molecular database (Zinc15, chEMBL, Asinex, Decoy molecules, ChemDiv, and Specs) sources.

**Scheme 1:**
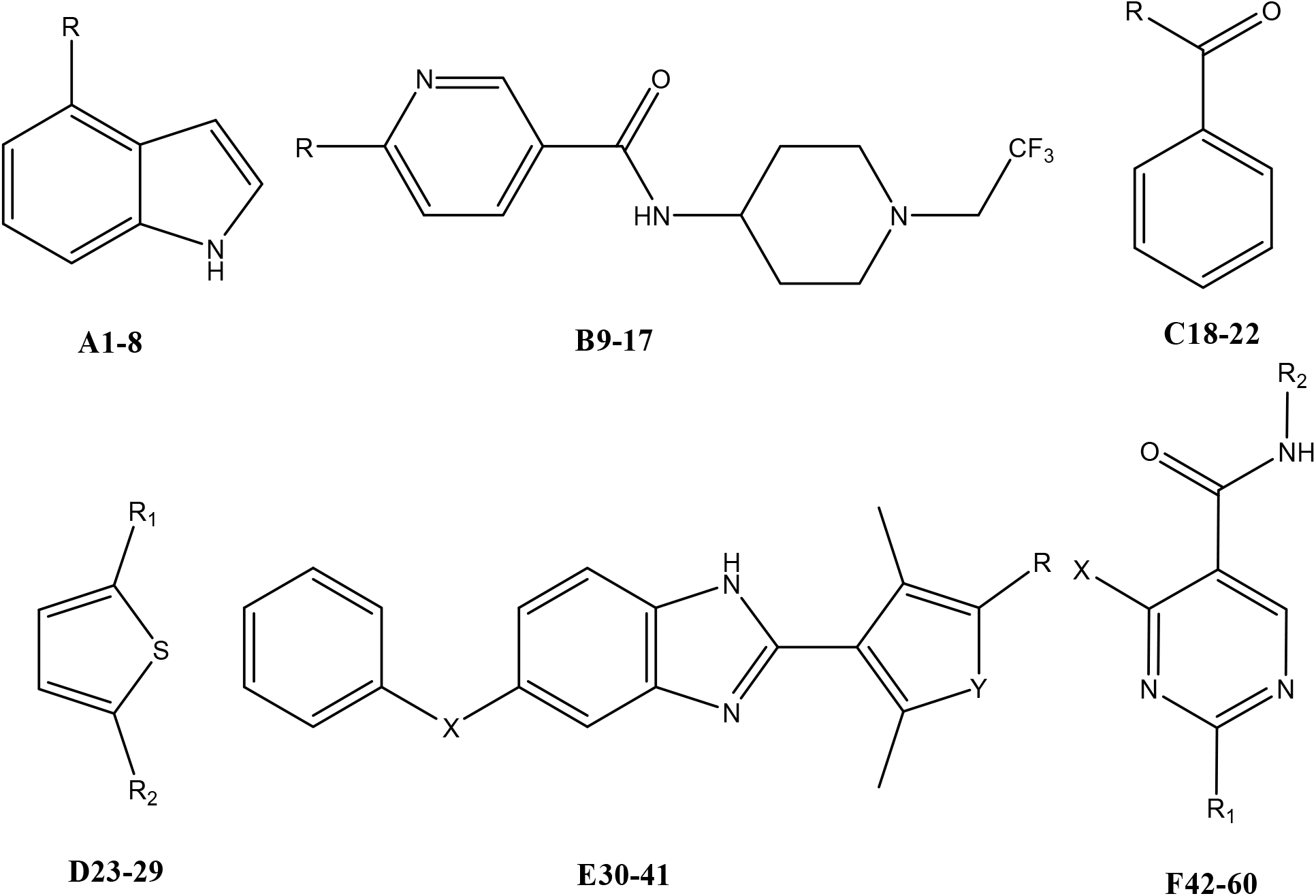
Chemical structure of molecules with indole **[A1-8]**, pyridine **[B9-17]**, benzaldehyde **[C18-22]**, thiophene **[D23-29]**, benzimidazole **[E30-41]** and pyrimidine **[F42-60]** scaffolds.

## 2. Methodology

Computational studies were performed using Maestro 11.9 module [37]. The process includes pharmacophore-based molecular screening, molecular docking, MM/GBSA, ADME analysis, and molecular dynamics (MD) simulations. The flowchart of the experimental work has been depicted in scheme 2.

**Scheme.2:**
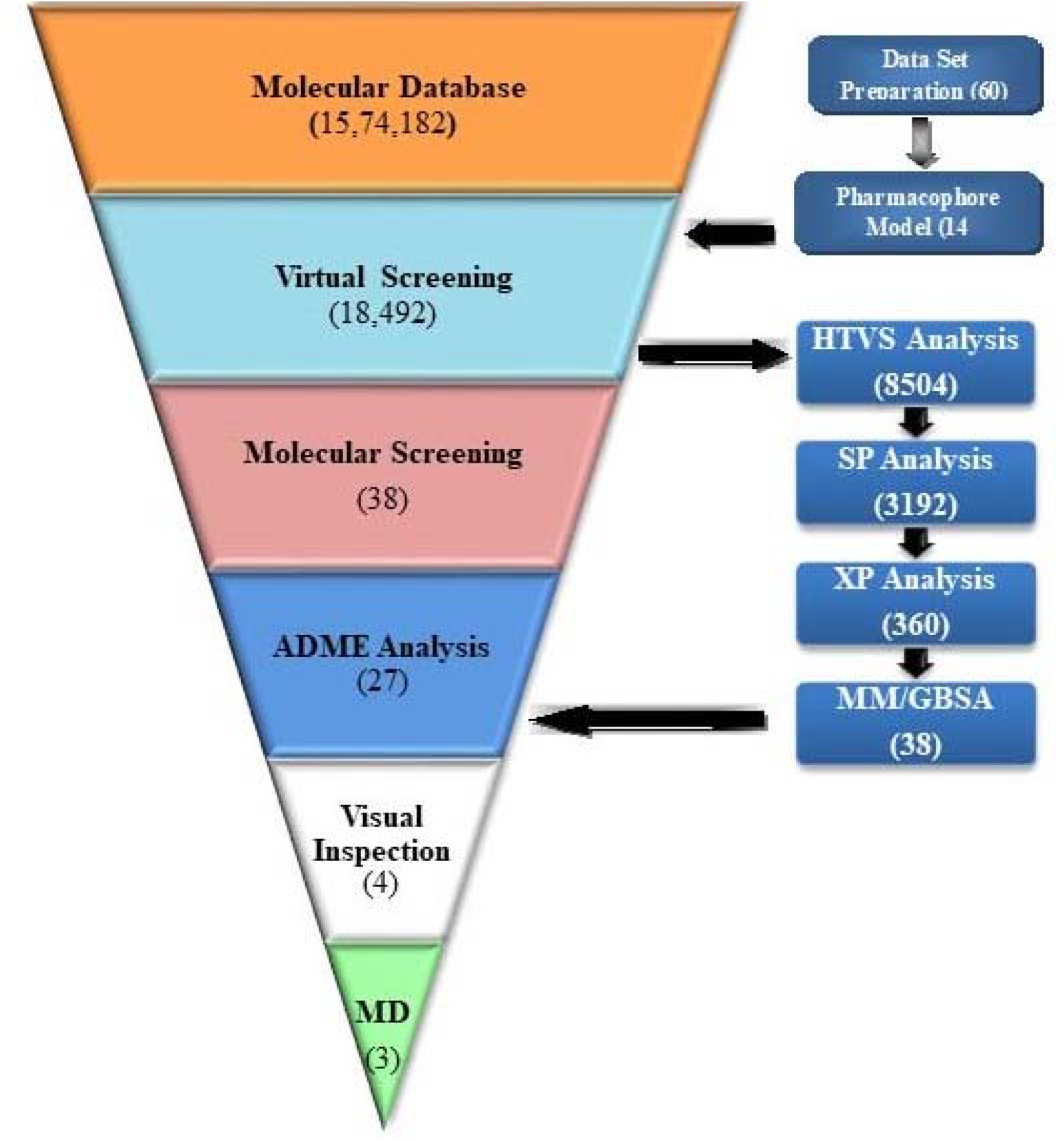
Working methodology

### 2.1 Dataset and Protein preparation

A set of 60 molecules based on a literature survey taken into consideration (**Table S1)**, having a spanned range of pIC_50_values (8.301 to 3.932).3D structures of all the molecules were generated in Maestro 11.9 and optimized in the ‘LigPrep’ module [37] by using the OPLS_2005 force field [38]. For structural optimization of the molecules, no tautomers were considered as well as only one stereoisomer (retaining specified chiralities) was generated per ligand and remaining were set as default.

3D crystal structure of the human HPGD2 enzyme (PDB ID-2CVD) [26] was downloaded from protein data bank having 1.45 Å resolution. Protein preparation is carried out by using the ‘Protein preparation wizard’ module. In the preprocessing of protein, missing hydrogens were added, deleted water molecules which are beyond 5 Å from hetero-group and ‘ionization and tautomeric’ states were generated at pH range (7.0 ± 2.0) by considering Epik (Empirical pKa Prediction). Hydrogen bonds were optimized at pH 7.0, as determined from a pKa prediction by PROPKA and finally, minimization was executed.

### 2.2 Pharmacophore based virtual screening

The pharmacophore model [39] was generated by using the ‘phase module’ of Schrodinger 2019-1. The optimized HPGD2 inhibitors were aligned by keeping the most bioactive molecule at the top as a template. The aligned molecules were used for the generation of a pharmacophore model. An activity threshold boundary of pIC50 > 7.49 and pIC50 < 5.43 were considered to generate a set 8 active, 8 inactive and 44 moderately active molecules. The best pharmacophore Hypothesis was selected based on BEDROC score, PhaseHypoScore, and Survival score.

A total of 15,74,182 molecules were collected from a different database (like Zinc15, chEMBL, Asinex, Decoy molecules, ChemDiv, and Specs) and bioactive HPGD2 inhibitors were screened by mapping them on the generated pharmacophore hypothesis. The virtual screening was performed by using the “Phase Ligand screening” tool of the Phase module. Screened 18,492 molecules were optimized and subjected to molecular screening through molecular docking and MM/GBSA study.

### 2.3 Molecular docking

Ligand docking was performed by using the ‘Ligand Docking’ module of Maestro 11.9.Prior to the operation, the active site was generated by using the grid generation module ‘Receptor Grid Generation’ by keeping van der Waals radius scaling factor 1.0 and partial charge cutoff at 0.25 and remaining set as default. For analyzing interaction behavior and binding affinity, all 18,492 molecules were docked in the active site of the minimized HPGD2 enzyme (PDB-ID:2CVD). Ligand docking was performed using different docking mode right from HTVS (High Throughput Virtual Screening) to SP (Standard Precision) to XP (Extra Precision), to screen the large number of 18,492 molecules on the basis of interaction behavior, binding affinity and docking score.

### 2.4 MM-GBSA

In Molecular Mechanics Generalized Born Surface Area (MM-GBSA), the binding free energy (ΔG_bind_) of the protein-ligand complex was calculated by using the prime ‘MM-GBSA module’. The energies of the protein-ligand complex were computed with an OPLS-2005 force field and VSGB (Energy Model for High-Resolution Protein Structure Modeling) [40] solvent model and remaining set as default. Molecules were screened on the basis of binding free energy (ΔG_bind_).

### 2.5 ADME Property analysis

Pharmacokinetics parameters and physicochemical properties of the molecules were calculated by using the ADME descriptors algorithm. ADME properties of all 38 molecules (based on MM-GBSA outcomes) were analyzed by using the ‘Qikprop’ module of Maestro 11.9 (Ligand-based ADME).Lipinski’s rule of five is used to evaluate drug-likeness and filter the molecules on the basis of druggability [41,42]. ADME explored the details of rule of five (like mol_MW< 500, QPlogPo/w < 5, donorHB ≤ 5, accptHB ≤ 10), predicted apparent Caco-2 cell permeability in nm/sec (**QPPCaco**) and predicted aqueous solubility (**QPlogS**).Molecules were screened out on the basis of violations of rules.

### 2.6 Molecular dynamics simulation

Molecular dynamics (MD) simulations were performed using Desmond from D. E. Shaw Research. The visual inspection outcome molecules i.e. **SP1**, **SP2**, **SP5**, **SP10** were docked with human HPGD2 enzyme (2CVD) and the docked file was further subjected to molecular dynamics simulations study. The protein was solvated in a three centered water model employing a simple point charge (SPC) solvent model in an orthorhombic box. Protein atoms were placed at a distance of 10 Å from the edge of the simulation box creating a buffer region between them. All 4 systems were neutralized by adding 5 Na+ ions and energy minimization was performed using the OPLS_2005 force field. LBFGS minimization was performed with three vectors and ten Steepest descent (SD) steps until a gradient threshold of 25 kcal/mol/Å was reached. The cutoff radius for short-range coulombic forces was 9.0 Å and the maximum iterations and convergence threshold was kept at 2000 & 1.0 kcal/mol/Å respectively. Further protein-ligand complex dynamics were simulated for 30 ns with the isotropic coupling of Nose-Hoover chain and Martyna-Tobias-Klein methods. In the NPT ensemble, the equilibrium phase was attained by heating the system to 300K at 1.01325 bar pressure with a relaxation time of 1ps employing both the before mentioned thermostat and barostat methods. RESPA integration algorithm for bonded (2.0 fs), near (2.0 fs) and far (6.0fs) were kept for multiple time step dynamics. After the simulation, the trajectories and 3D structures were inspected from the generated simulation interaction diagram.

## 3. Result and Discussion

### 3.1 Pharmacophore based virtual screening

The Pharmacophore Hypothesis generated 14 models (table 1) with 6 different types like ADRR (model 1, 5 and 10), AADR (model 2, 3, 4 and 7), AAADR (model 6 and 8), AADRR (model 13 and 14), AAAR (model 9) and AAAD (model 11 and 12). The four featured pharmacophore model ADRR_1 considered the best model as it shows the best BEDROC score (0.839), best PhaseHypoScore (1.110) and good Survival Score (4.518). ADRR_1 consists of one H-bond acceptor (red color), one H-bond donor (blue color)and two Aromatic rings (orange color) (fig.2a and the position of features concerning each other shown in fig.2b). ADRR_1 Hypothesis shows eight active molecules (**F-42**, **F-55**, **F-47**, **F-44**, **F-43**, **B-11**, **A-5** and **A-4**) and two inactive molecules (**C-21** and **C-22**). **F-42 (**pIC_50_-8.301),the most bioactive molecule consists of all the features of ADRR_1 model whereas the inactive **C-22** (pIC_50_-3.932), features did not match with the hypothesis. The molecules screened from various databases were mapped on the best pharmacophore model ADRR_1 and collected 18,492 molecules. This Pharmacophore based virtual screening symbolize that 18,492 molecules had the probability to bind the target protein out of 15,74,182 molecules. A primary elimination procedure was completed through the pharmacophore-based virtual screening.

**Table 1:**
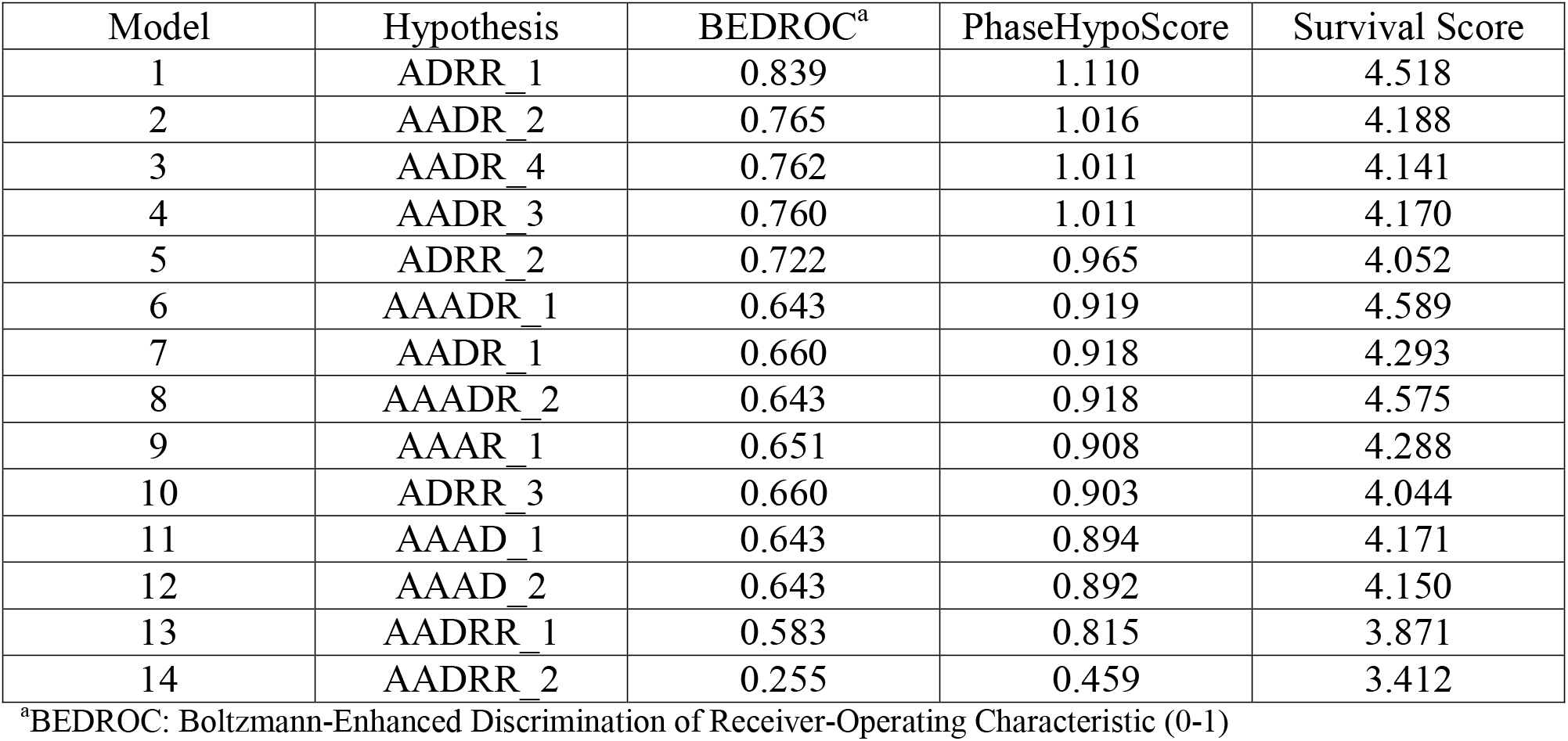
Phase generated pharmacophore hypothesis for the hematopoietic prostaglandin D2 synthaseinhibitor.

**Fig.2a:**
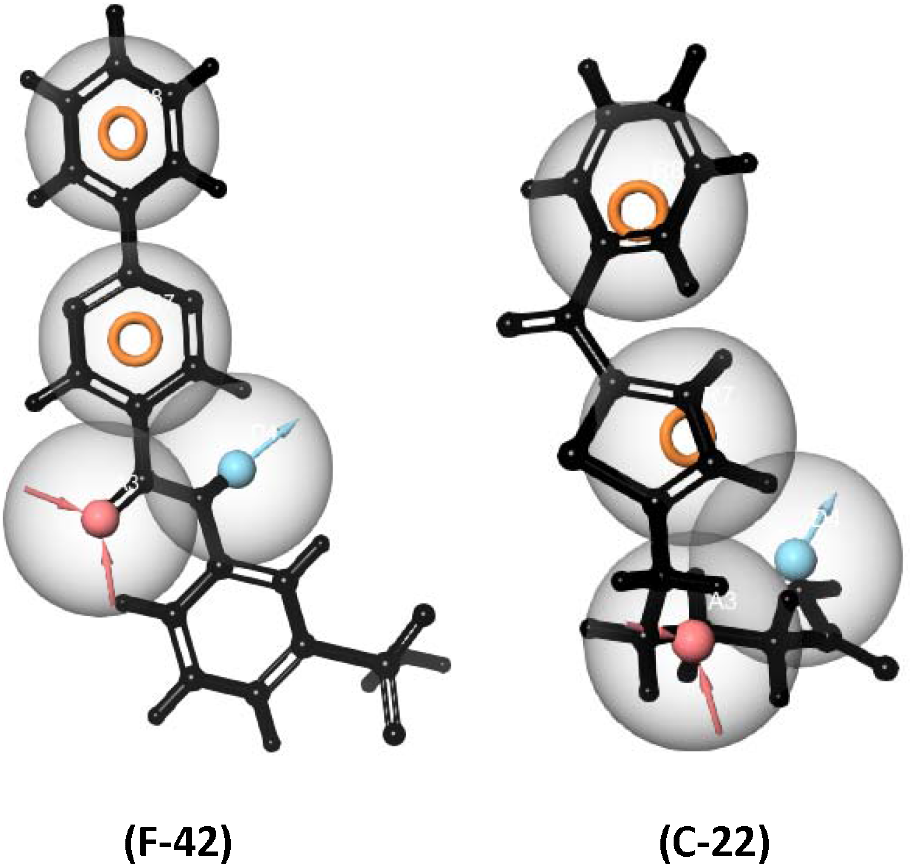
Mapping of molecules (a1) active molecule F-42 (pIC_50_-8.301) and (a2) inactive molecule C-22 (pIC_50_-4.517) on pharmacophore hypothesis model ADRR_1 for HPGD2 inhibitor.

**Fig.2b:**
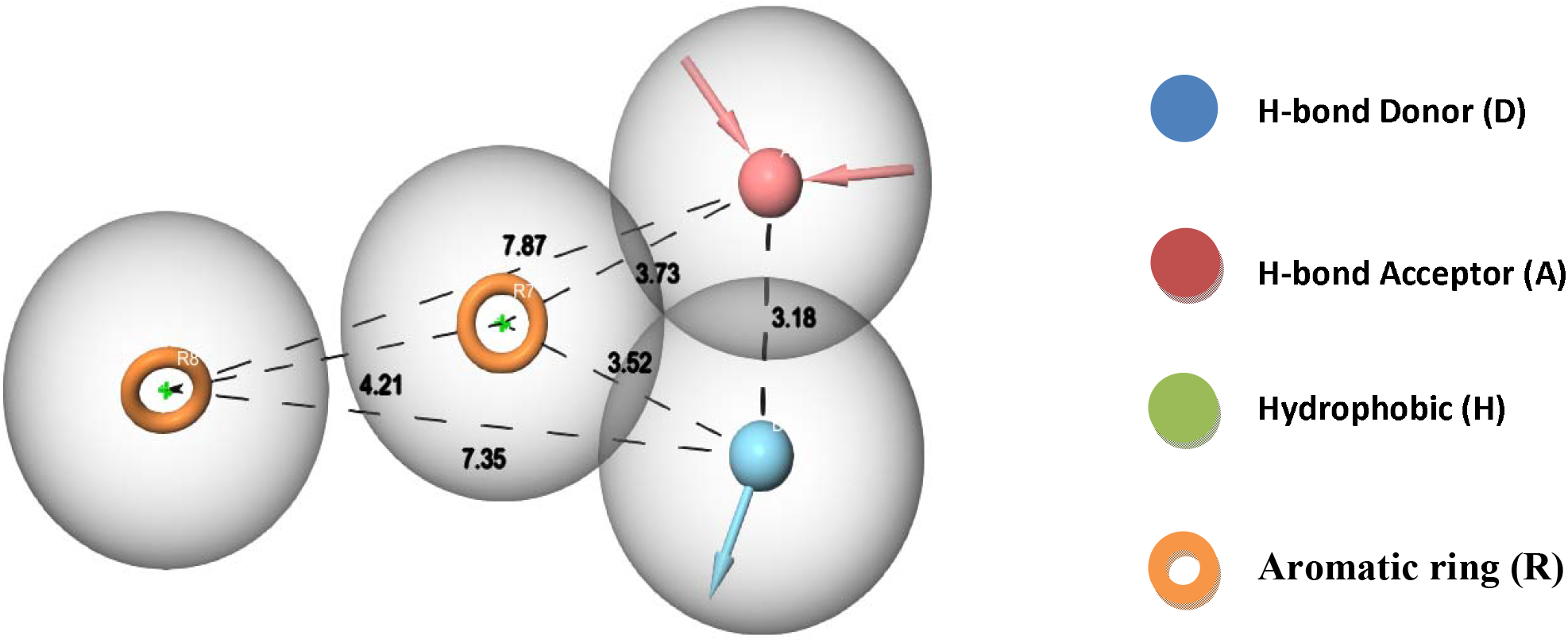
Arrangement of individual features in a fixed distance (in A^0^) of pharmacophore hypothesis model ADRR_1 for HPGD2 inhibitor.

### 3.2 Molecular docking

The generated active site of human hPGDS (PDB_ID: 2CVD) is occupied by charged amino acids (ARG-12, ARG-14, LYS-50 and ASP-96), polar amino acids (SER-100 and THR-159), hydrophobic amino acids (TYR-8, PHE-9, MET-11, IEL-51, IEL-55, MET-99, PHE-102, TRP-104, TYR-150, ILE-155, CYS-156, LUE-160, LUE-199) and one glycine (GLY-13) residue. To ensure the excavation of the best candidates, molecular screening was performed through ligand docking into the active site of the hPGDS enzyme. Due to the huge data set, preliminary screening was done by HTVS (High-Throughput Virtual Screening) method. This method reduced the active molecule count from 18,492 to 8,504. Preliminary screened molecules were subjected to SP (Standard Precision) mode, which further reduced the molecule count to 3,192. The final screening of molecular docking was performed by XP (Extra Precision) mode and the resultant set of 360 molecules was generated. The various molecular docking modes at each step of ligand docking increases the accuracy to predict molecule-protein interactions.

### 3.3 MM-GBSA

After the screening of a huge database through molecular docking, the resultant molecules (360) were subjected to ligand-receptor binding energy, MM-GBSA analysis. The MM-GBSA analysis computed the binding free energy of docked ligand-receptor complex which confirms the stability of the ligand after binding to the active site of the enzyme. The generated ‘MM-GBSA_dG_Bind’ energy by using the MM-GBSA module shows the energy difference between prime energy (optimized ligand-receptor complex) and the combined energy of optimized free ligand and optimized free receptor. Out of 360 molecules, 38 were selected based on the MM-GBSA_dG_Bind energies i.e. more than 60 kcal/mole was taken into consideration. The MM-GBSA_dG_Bind energies for all 38compounds are shown in Table 2.

**Table 2:** MM-GBSA_dG_Bindvalues of 38screened molecules in (kcal/mol) obtained through MM-GBSA analysis.

| No<br>. | title | MMGBSA_dG_Bind | No<br>. | title | MMGBSA_dG_Bind |
| --- | --- | --- | --- | --- | --- |
| 1 | SP1 | -62.20626292 | 20 | SP20 | -63.90098569 |
| 2 | SP2 | -64.71327396 | 21 | SP21 | -65.00114709 |
| 3 | SP3 | -64.35759475 | 22 | SP22 | -61.8611676 |
| 4 | SP4 | -60.48284789 | 23 | SP23 | -62.96275558 |
| 5 | SP5 | -67.23541626 | 24 | SP24 | -61.8441487 |
| 6 | SP6 | -62.60572487 | 25 | SP25 | -63.82207561 |
| 7 | SP7 | -64.26938416 | 26 | SP26 | -63.55243427 |
| 8 | SP8 | -60.52766668 | 27 | SP27 | -61.27199217 |
| 9 | SP9 | -65.14678131 | 28 | SP28 | -65.7207912 |
| 10 | SP10 | -61.5618593 | 29 | SP29 | -61.53606989 |
| 11 | SP11 | -63.55415521 | 30 | SP30 | -61.13560865 |
| 12 | SP12 | -63.32152295 | 31 | SP31 | -61.16275959 |
| 13 | SP13 | -61.08042816 | 32 | SP32 | -62.18381642 |
| 14 | SP14 | -60.50102233 | 33 | SP33 | -63.35208723 |
| 15 | SP15 | -63.49689636 | 34 | SP34 | -63.24818574 |
| 16 | SP16 | -63.58435786 | 35 | SP35 | -67.19227179 |
| 17 | SP17 | -60.01429792 | 36 | SP36 | -64.33664826 |
| 18 | SP18 | -61.78569113 | 37 | SP37 | -62.7408134 |
| 19 | SP19 | -61.00111018 | 38 | SP38 | -60.60995987 |

### 3.4 ADME analysis

The drug-likeness properties of the molecules studied by using well-known ADME analysis i.e., Absorption (A), Distribution (D), Metabolism (M) and Excretion (E) which explains the disposition of pharmaceutical compound inside an organism and therefore, influences the pharmacological activity of it. The screening was performed based on a violation of ‘Lipinski’s rule of five ‘, **QPPCaco** and **QPlogS**. In the ADME property analysis out of 38 molecules, 27 were shown no violation of ‘Lipinski’s rule of five’, (**QPPCaco**) and (**QPlogS**) (Table 3) whereas remaining 11 molecules were violating above parameters (shown in the bold in Table 3). (detailed structures are shown in Table S2).

**Table 3:** ADME properties of all 38 ligands to determine their ‘drug-likeness’.

| molecule | mol_MW | donorHB | accptHB | QPlogPo/w | RuleOfFive | QPPCaco | QPlogS |
| --- | --- | --- | --- | --- | --- | --- | --- |
| SP1 | 295.297 | 3 | 10.4 | -0.392 | 0 | 60.981 | -2.463 |
| SP2 | 265.271 | 4 | 8.9 | -0.206 | 0 | 122.838 | -2.099 |
| **SP3** | **295.303** | **2** | **11.2** | **-1.515** | **1** | **10.477** | **-0.336** |
| **SP4** | **294.316** | **4** | **9.7** | **-1.007** | **0** | **14.91** | **-1.022** |
| SP5 | 281.271 | 3 | 9.9 | -0.457 | 0 | 81.984 | -2.108 |
| **SP6** | **296.303** | **3.25** | **9.45** | **-1.299** | **0** | **14.342** | **-1.946** |
| SP7 | 281.271 | 3 | 10.4 | -0.717 | 0 | 55.562 | -2.165 |
| **SP8** | **293.285** | **2** | **8.5** | **-0.585** | **0** | **17.686** | **-1.904** |
| SP9 | 296.285 | 3 | 11.9 | -1.571 | 0 | 60.69 | -1.441 |
| SP10 | 293.285 | 4 | 10.7 | -1.027 | 0 | 102.39 | -0.791 |
| SP11 | 280.286 | 3 | 9.9 | -0.372 | 0 | 67.826 | -2.399 |
| SP12 | 291.266 | 0.25 | 7.45 | 0.549 | 0 | 58.265 | -2.701 |
| SP13 | 278.316 | 1 | 8 | -0.42 | 0 | 30.577 | 0.186 |
| SP14 | 293.331 | 0 | 10 | -1.019 | 0 | 38.261 | 0.458 |
| SP15 | 292.3 | 3 | 10 | -0.831 | 0 | 33.082 | -2.005 |
| **SP16** | **294.316** | **3** | **9.2** | **-1.01** | **0** | **9.937** | **-0.493** |
| SP17 | 247.256 | 2 | 7.7 | 0.019 | 0 | 146.566 | -1.374 |
| SP18 | 281.274 | 3 | 10.9 | -0.861 | 0 | 110.468 | -1.612 |
| SP19 | 292.3 | 2 | 9 | 0.166 | 0 | 62.632 | -3.397 |
| SP20 | 296.303 | 3 | 9.7 | -0.712 | 0 | 33.336 | -1.827 |
| SP21 | 241.267 | 1 | 8.4 | 0.177 | 0 | 635.793 | -1.288 |
| SP22 | 278.27 | 2 | 9.9 | -0.388 | 0 | 103.187 | -1.878 |
| SP23 | 294.316 | 2 | 8.5 | -0.08 | 0 | 26.558 | -1.724 |
| SP24 | 237.261 | 1 | 7.4 | 0.777 | 0 | 736.711 | -1.461 |
| **SP25** | **295.303** | **3** | **13** | **-2.055** | **1** | **13.036** | **-0.938** |
| SP26 | 294.313 | 2 | 9.4 | -0.19 | 0 | 142.156 | -0.743 |
| SP27 | 292.3 | 2 | 9 | -0.459 | 0 | 39.152 | -2.006 |
| **SP28** | **319.325** | **0** | **10** | **-1.541** | **1** | **11.567** | **-0.213** |
| SP29 | 291.312 | 3 | 7 | 1.128 | 0 | 111.081 | -3.828 |
| **SP30** | **293.331** | **2** | **10** | **-0.961** | **0** | **17.258** | **-0.836** |
| **SP31** | **293.331** | **3** | **9.5** | **-0.626** | **0** | **14.36** | **-1.623** |
| SP32 | 293.331 | 0 | 9.5 | -0.889 | 0 | 35.049 | 0.39 |
| SP33 | 266.259 | 4 | 9.9 | -1.139 | 0 | 37.347 | -1.861 |
| **SP34** | **291.315** | **1** | **9** | **-0.592** | **0** | **16.833** | **-0.844** |
| SP35 | 291.315 | 1 | 9 | -0.438 | 0 | 37.387 | -0.73 |
| **SP36** | **291.315** | **2** | **10** | **-1.144** | **0** | **9.426** | **-0.726** |
| SP37 | 290.284 | 2 | 9 | -0.075 | 0 | 98.605 | -2.199 |
| SP38 | 265.274 | 1 | 9.2 | 0.196 | 0 | 427.122 | -1.875 |

### 3.5 Visual Inspection

The selected 27 molecules (Table 4) were inspected individually based on the docking score, binding free energy and ligand-receptor interaction diagram, the best 4 structures were selected i.e. **SP1, SP2, SP5, SP10**. The selected molecules were showing docking score and binding free energy like −8.41, −7.79, −6.16, −5.89 and −62.20626292 kcal/mol, −64.71327396 kcal/mol, −67.23541626 kcal/mol, −61.5618593 kcal/mol respectively. Ligand **SP1** shows 3 H-bonds with hydrophobic amino acids (ILE 51, TRP 104), two H-bonds due to H-bond donor group and one due to H-bond acceptor site present in the ligand.(Fig.3) **SP2** shows 4 H-bond interactions due to two H-bond donor groups and two H-bond acceptor groups present in the ligand as well as ligand also shows two pi-pi stacking interactions. All the interactions take place with the hydrophobic amino acid present in the active site.(Fig.3) **SP5** shows 4 H-bond interactions due to 3 H-bond donor groups and one H-bond acceptor group present in the ligand, it also shows two pi-pi stacking interactions and all the interactions take place with the hydrophobic amino acid present in the active site.(Fig.3) **SP10** shows 5 H-bond interactions due to 2 H-bond donor groups and 3 H-bond acceptor groups present in the ligand. It also shows three pi-pi stacking and one pi-cation interaction.(Fig.3) All these interactions take place with the hydrophobic amino acid present in the active site. All 4 ligands were showing a good range of solvent exposure.

**Table 4:** Docking score and functional parameters of all 27 visually inspected molecules and commercially available compounds used in this study.

| ligand | Dock Score | LipophilicEvdW | PhobEn | PhobEnHB | HBond | Electro | LowMW |
| --- | --- | --- | --- | --- | --- | --- | --- |
| **SP1** | -8.41 | -2.45 | -1.39 | -1 | -3.4 | -0.7 | -0.5 |
| **SP2** | -7.79 | -2.75 | 0 | -1.5 | -3.2 | -0.98 | -0.5 |
| **SP5** | -6.16 | -2.72 | -1.2 | 0 | -2.19 | -0.74 | -0.5 |
| SP7 | -6.09 | -2.67 | -0.85 | 0 | -2.19 | -0.7 | -0.5 |
| SP9 | -6.07 | -2.95 | 0 | 0 | -3.05 | -1 | -0.5 |
| **SP10** | -5.89 | -3.1 | 0 | 0 | -2.26 | -0.89 | -0.5 |
| SP11 | -5.88 | -2.14 | 0 | -0.83 | -2.85 | -0.94 | -0.5 |
| SP12 | -5.72 | -2.93 | -0.2 | 0 | -1.92 | -0.6 | -0.5 |
| SP13 | -5.37 | -3.27 | 0 | -1.5 | -0.73 | -0.32 | -0.5 |
| SP14 | -5.32 | -2.85 | -0.9 | 0 | -1.63 | -0.41 | -0.5 |
| SP15 | -5.3 | -2.65 | 0 | -1.5 | -1.01 | -0.5 | -0.5 |
| SP17 | -5.18 | -3.01 | -2.02 | 0 | -0.47 | -0.19 | -0.5 |
| SP18 | -5.13 | -3.07 | 0 | 0 | -1.33 | -0.87 | -0.5 |
| SP19 | -5.09 | -2.72 | 0 | 0 | -2.08 | -0.55 | -0.5 |
| SP20 | -5.04 | -3.19 | 0 | 0 | -1.48 | -0.41 | -0.5 |
| SP21 | -5.03 | -3.11 | -0.9 | 0 | -1.49 | -0.31 | -0.5 |
| SP22 | -4.92 | -3.27 | 0 | 0 | -1.92 | -0.48 | -0.5 |
| SP23 | -4.84 | -3.63 | 0 | 0 | -0.7 | -0.31 | -0.5 |
| SP24 | -4.83 | -3.85 | 0 | 0 | -1.74 | -0.57 | -0.5 |
| SP26 | -4.56 | -3.19 | 0 | 0 | -1.24 | -0.68 | -0.5 |
| SP27 | -4.36 | -2.67 | -0.55 | 0 | -1.34 | -0.52 | -0.5 |
| SP29 | -3.89 | -2.7 | 0 | 0 | -1.23 | -0.57 | -0.5 |
| SP32 | -3.88 | -3.15 | 0 | 0 | -0.7 | -0.31 | -0.5 |
| SP33 | -3.86 | -3.52 | 0 | 0 | -1.33 | -0.63 | -0.5 |
| SP35 | -3.67 | -2.99 | 0 | 0 | -0.56 | -0.23 | -0.5 |
| SP37 | -3.22 | -2.3 | 0 | 0 | -0.62 | -0.52 | -0.5 |
| SP38 | -3.18 | -2.54 | 0 | 0 | -1.05 | -0.38 | -0.5 |
| *TFC-007* | -5.3 | -4.1 | -0.2 | 0.0 | -0.7 | -0.3 | 0.0 |
| *HPGDS inhibitor I* | -6.01 | -3.1 | -2.1 | 0.0 | -0.6 | -0.3 | -0.2 |
| *HQL-79* | -6.09 | -3.9 | -1.8 | 0.0 | -0.7 | -0.4 | -0.2 |
| *TAS-204* | -6.11 | -3.9 | -1.6 | 0.0 | -0.7 | -0.23 | -0.1 |

**Fig.3:**
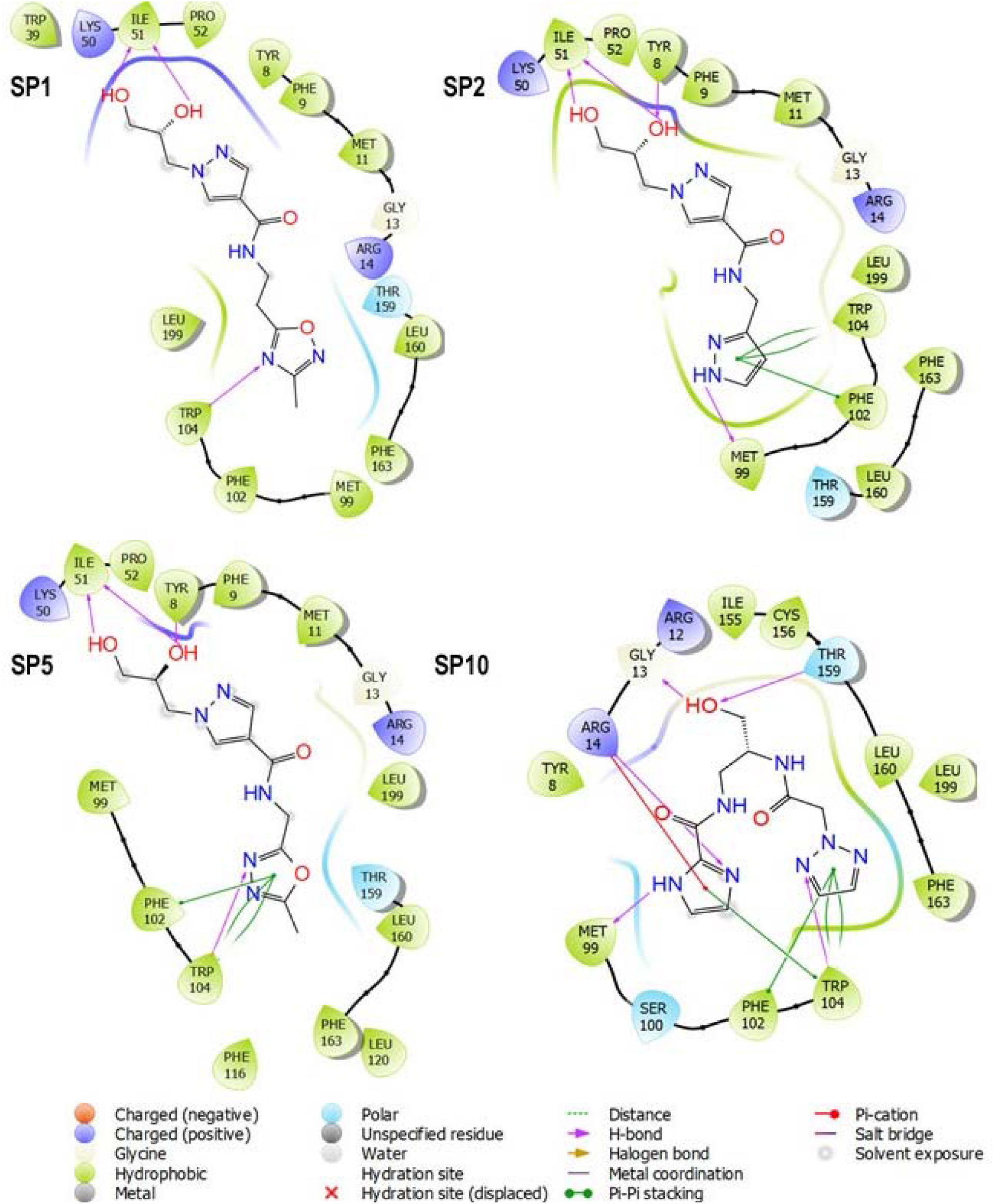
Molecular docking of active **SP1, SP2, SP5** and **SP10** molecules with human hematopoietic prostaglandin D2 synthase enzyme.

### 3.6 Comparative study

The comparative analysis of marketed compounds with screened molecules was studied. The marketed compounds like TFC-007, HPGDS inhibitor I, HQL-79 and TAS-204 were showing docking score −5.3, −6.01, −6.09 and −6.1 respectively. (Table 4) TFC-007 [27] showing two H-bond, hPGDS inhibitor-I [28] showing one H-bond and two pi-pi stacking, HQL-79 [24–26] showing two H-bond and one pi-pi stacking and TAS-204 [29] also showing two H-bond and one pi-pi stacking. (Fig.4) The comparative study concludes that the screened compounds showing better interaction in comparison to marketed compounds.

**Fig.4a:**
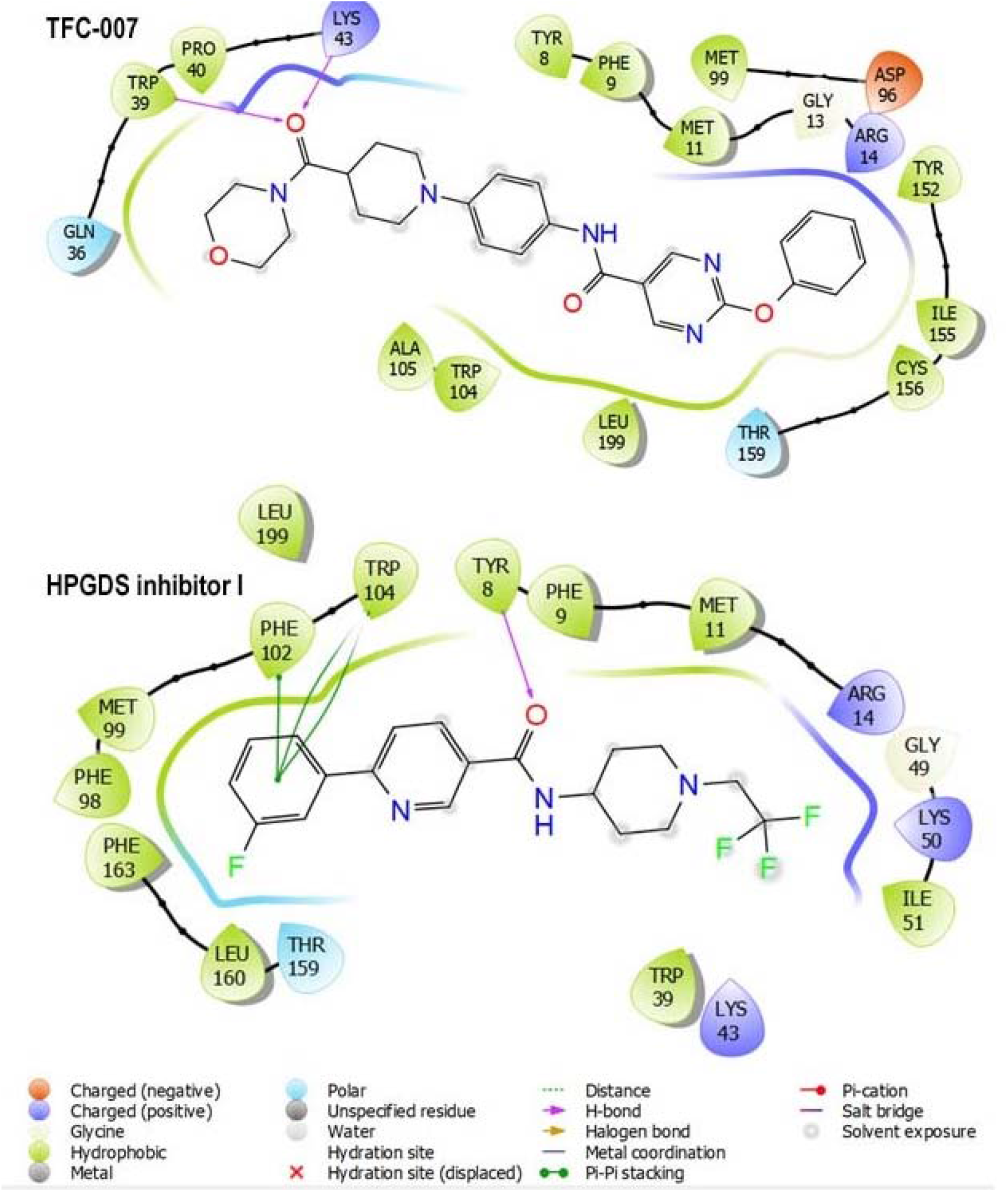
Molecular docking interactions of commercially available (TFC-007 and HPGDS inhibitor I) compounds with human hematopoietic prostaglandin D2 synthase enzyme.

**Fig.4b:**
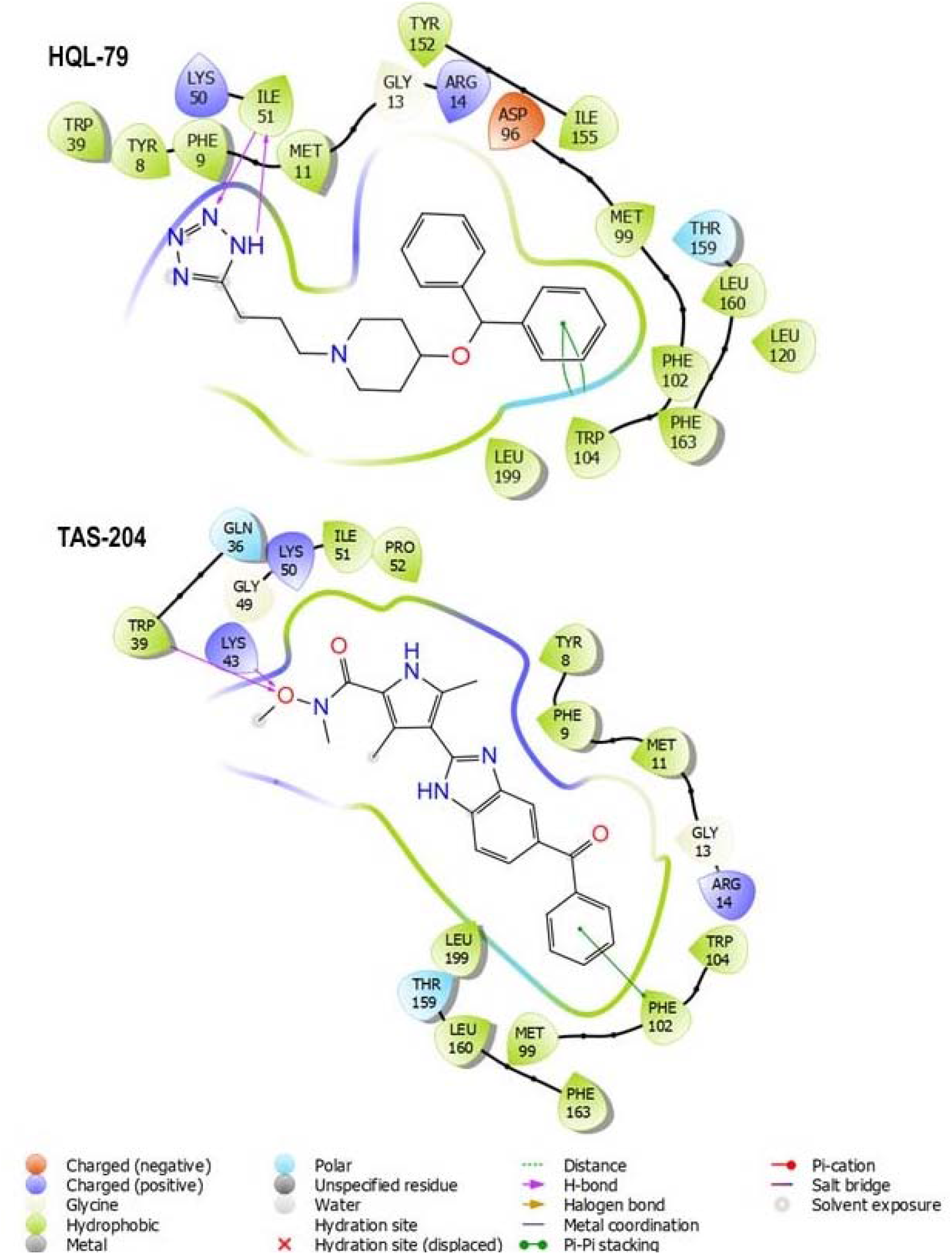
Molecular docking interactions of commercially available(HQL-79 and TAS-204)compounds with human hematopoietic prostaglandin D2 synthase enzyme.

### 3.7 Molecular dynamics simulations

MD simulations were performed to study the physical movements of atoms & molecules and the dynamic evolution of the entire system. The RMSD is a quantitative parameter to estimate the stability of the protein-ligand system.(Fig.5) The RMSD trajectory of Human PGD2 (**2CVD**) and ligand (**SP1, SP2, SP5, SP10)** complex shows a heavy fluctuation up to 24 ns simulation time then gradually tends to equilibrium. The RMSD average value of **2CVD-SP1, 2CVD-SP2, 2CVD-SP5** and **2CVD-SP10** complexes after reaching equilibrium were 2.0 □, 1.8 □, 3.2□, and 1.75□ respectively.(Fig.5) The RMSD curve of **2CVD-SP1, 2CVD-SP2** and **2CVD-10** is more stable in comparison with **2CVD-SP5** as for the small globular protein the deviation within 1-3 Å is acceptable. Based on the stability, **2CVD-SP5** was excluded and ‘**2CVD-SP1, 2CVD-SP2** and **2CVD-SP10’** were considered for further study like Protein-Ligand contact study (Histograms) and Ligand-Protein Contact study (ligand-receptor interaction diagram).

**Fig.5:**
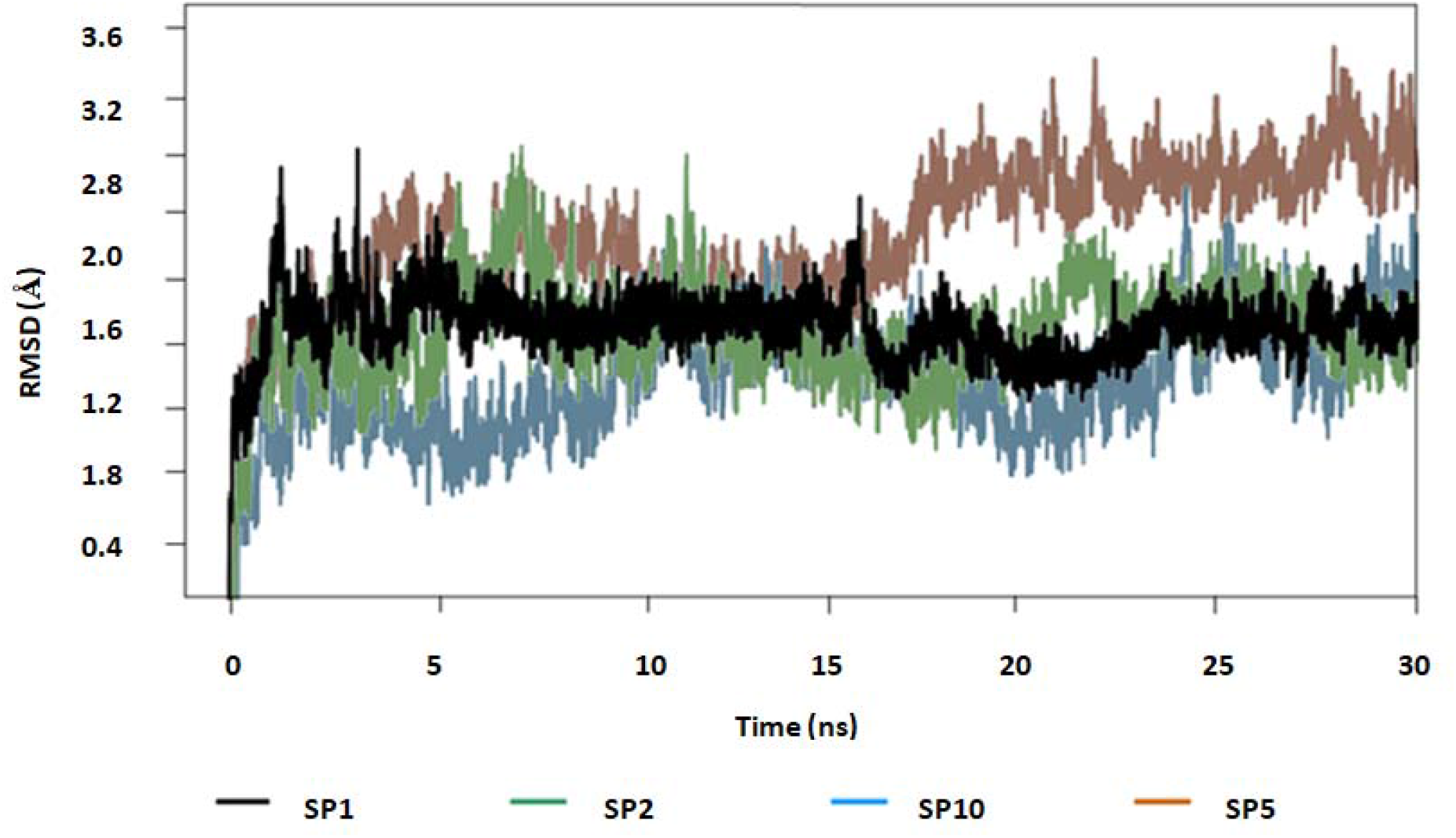
The RMSD trajectory of the human PGD2-ligand complex during the 30 ns simulation.

The protein-ligand contacts of the stable **2CVD-SP1, 2CVD-SP2** and **2CVD-SP10** ligand-receptor complex were studied by using the protein-ligand contact histograms.(Fig.6) In the entire three complexes, respective ligands show some similar H-bonding with the protein i.e. TRP-8, ARG-14, SER-100, and TRP-104. **SP1** shows interactions with ARG-12, GLN-36, TRP-39, ILE-51, THR-159, LYS-50, and LYS-198 residues whereas **SP2** and **SP10** show interaction with’GLN-49, ILE-51, ASP-96, and MET-99’ and ‘ARG-12, GLN-36, PHE-102, THR-158, THR-159’ amino acid residue respectively. The histogram also shows a wide range of hydrophobic interactions in all the protein-ligand contacts. Some similar hydrophobic interactions were also seen in all three complexes with PHE-9, MET-11, ARG-14, MET-99 and TRP-104 amino acid residues. Additionally, **SP1** shows interactions with TRP-39, ALA-105, PHE-116, LEU-160, PHE-163, LEU-199 amino acid residues whereas **SP2** and **SP10** show interaction with ‘TRP-39’ and ‘PHE-102, ILE-155, VAL-162, PHE-163, LEU-199’ amino acid residues. Along with H-bond and hydrophobic interactions, the ligand-protein complex also shows water bridge as well as ionic interactions. The water bridge interactions were formed with almost all major interacting amino acids in all three ligand-protein complexes (fig.6). All three ligand-receptor complexes were showing very minimal ionic interaction as shown in fig.6.

**Fig.6:**
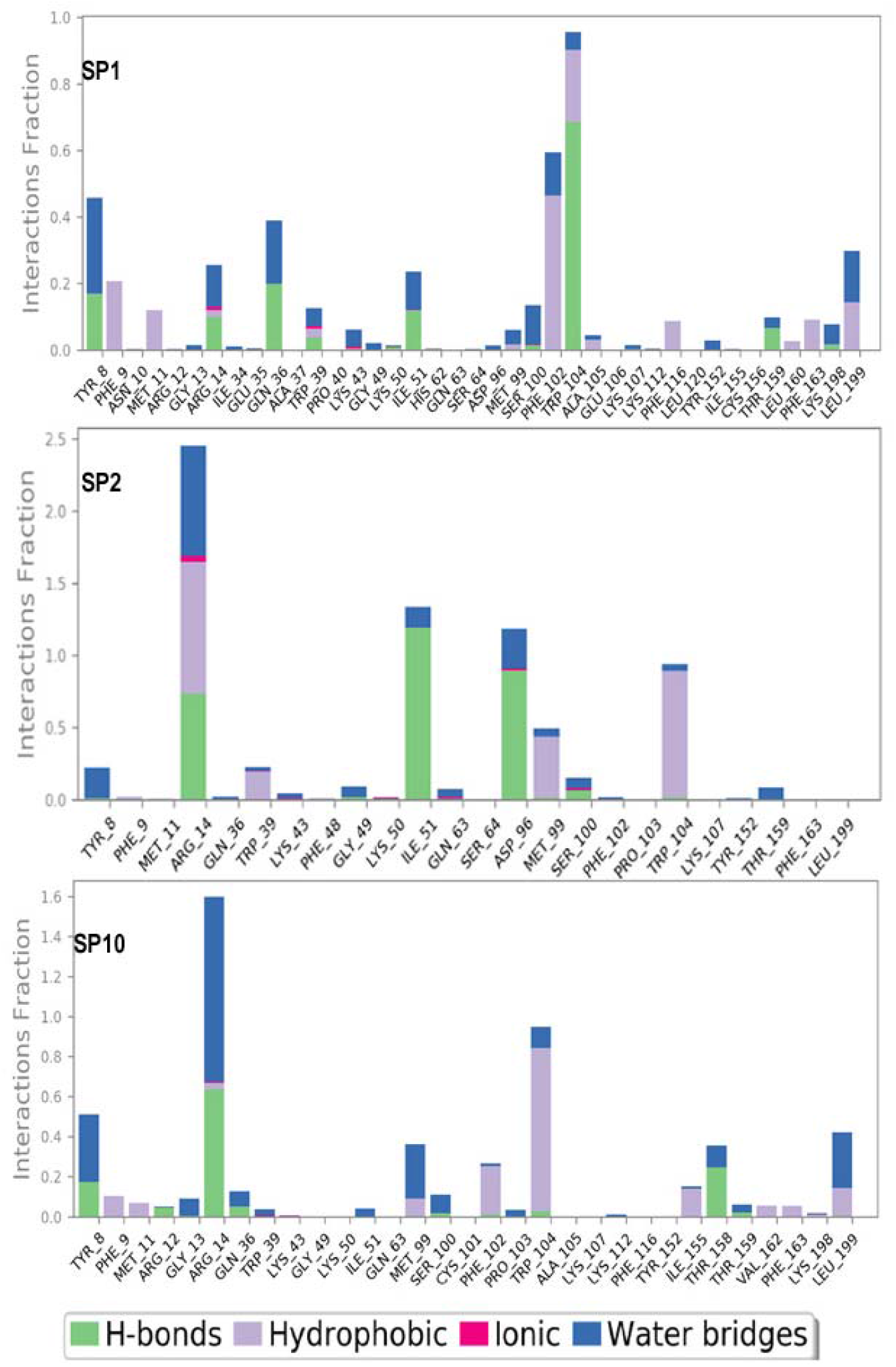
The histogram of protein-ligand contact over the course of the trajectory.

In protein-ligand contact histograms some amino acids were showing highly effective interactions like ‘PHE-102 and TRP-104’ having 61% and 95% interactions in **2CVD-SP1** complex, ARG-14, ILE-51, ASP-96 and TRP-104 having 240%, 140%, 130% and 94% interactions respectively in **2CVD-SP2** complex and ARG-14 and TRP-104 having 150% and 94% interactions respectively in **2CVD-SP10** complex.

The 30 ns MD simulation was divided into 6250 total number of trajectory frames and a single trajectory frame takes 4.8 ps time. The amino acid residues interact with the ligand in each trajectory frame shown in fig.S1. The repeated number of small lines in a band of amino acid row (fig.S1) represents all possible ligand-receptor interactions in 6250 trajectory frames. The **2CVD-SP1** receptor-ligand complex shows two deep bands (PHE-102 and TRP-104 row), **2CVD-SP2** shows four deep bands (ARG-14, ILE-51, ASP-96 and TRP-104 row) and **2CVD-SP10** complex shows two deep bands (ARG-14 and TRP-104 row) which explain that the above amino acid have more interaction with the ligands in almost all possible orientations (geometry) which is exactly similar as histogram results.

Ligand-Protein contacts were explored by using the ligand-receptor diagram generated by MD simulation.(Fig.7) For this study amino acid-ligand interactions, more than 4% were considered. Ligand **SP1** shows three pi-pi stacking interactions withPHE-102, TRP-104, and PHE-9 with a range of 17%, 8%, 7%. Ligand also shows six water bridges and eight hydrogen bond interactions (fig.7). Majorly TRP-104 shows an interaction of 67% with the nitrogen atom of 1,2, 4-oxadiazole in the ligand. Ligand **SP2** shows one pi-pi stacking interaction with TRP-104 at 40% and two pi-cationic interactions with ARG-14 at 47% and 44%. Ligand also shows sixteen water bridges and seven hydrogen bonds (fig.7). Some amino acids also shows major water bridge interactions like ARG-14 show four interactions with 36%, 17%, 10%, 4% and ASP-96 shows the interaction of 23% with the ligand. H-bonding major interactions were shown by ILE-51(65%, 43%, 9%), ARG-14 (57%, 15%), ASP-96 (88%) with the ligand. **SP10** ligand shows three pi-pi stacking interactions with PHE-102 (6%) and TRP-104 (21% and 8%). Ligand also shows sixteen water bridges and six hydrogen bonds (fig.7). Some amino acid shows major water bridge interaction like ARG-14 shows four interactions of ‘33%, 31% and 8%’ and TYR-8 shows interaction ‘21% and 6%’ with the ligand. H-bonding major interactions were shown by ARG-14 (36%, 11%), THR-158 (24%), TYR-8 (12%, 4%) with the ligand. All three ligands were showing solvent exposures but ligand **SP1** shows better exposures. The MD simulation study concludes that **SP1**, **SP2** and **SP10** ligands were more stable and having the best ligand-protein interactions out of 15,74,182 molecules.

**Fig.7:**
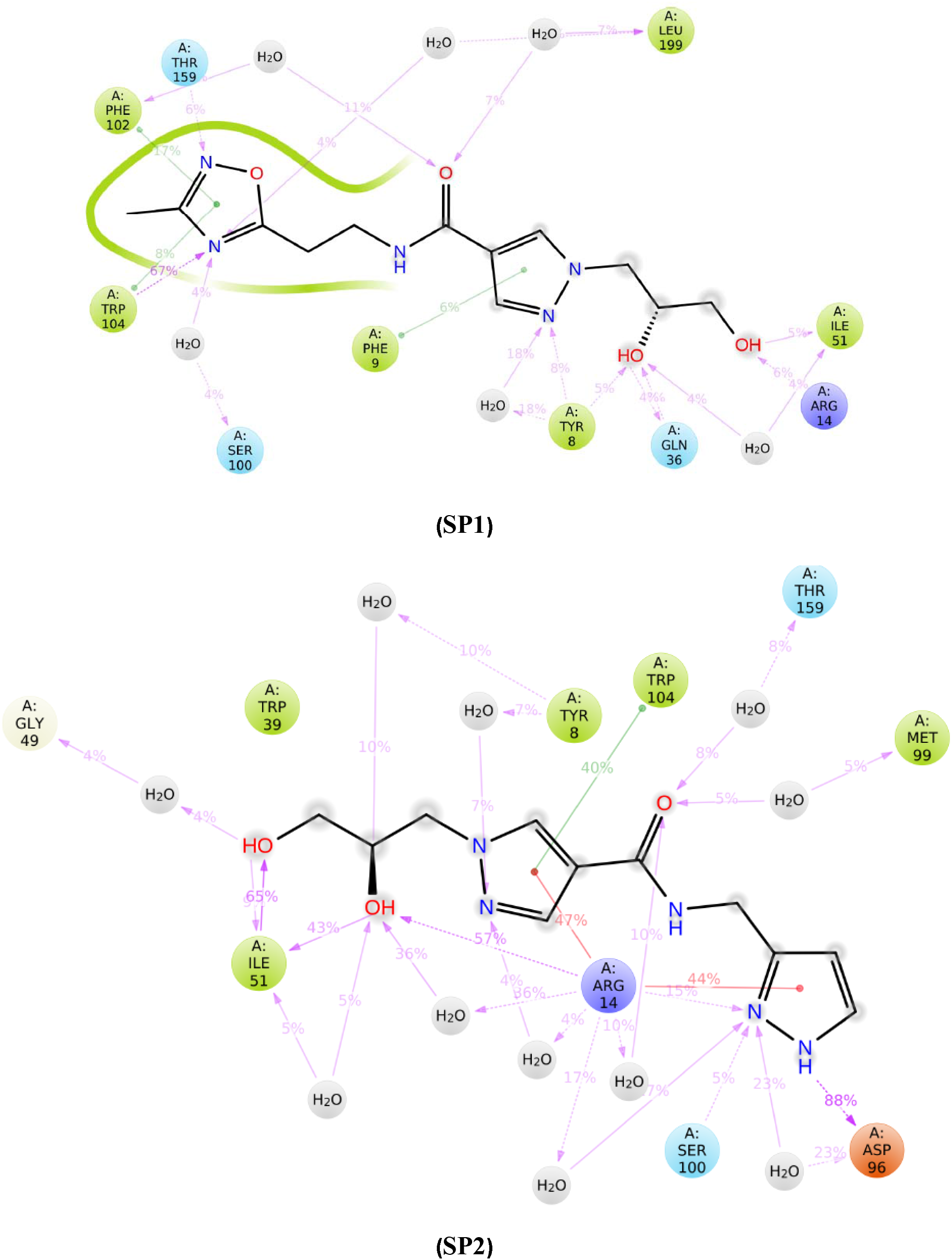
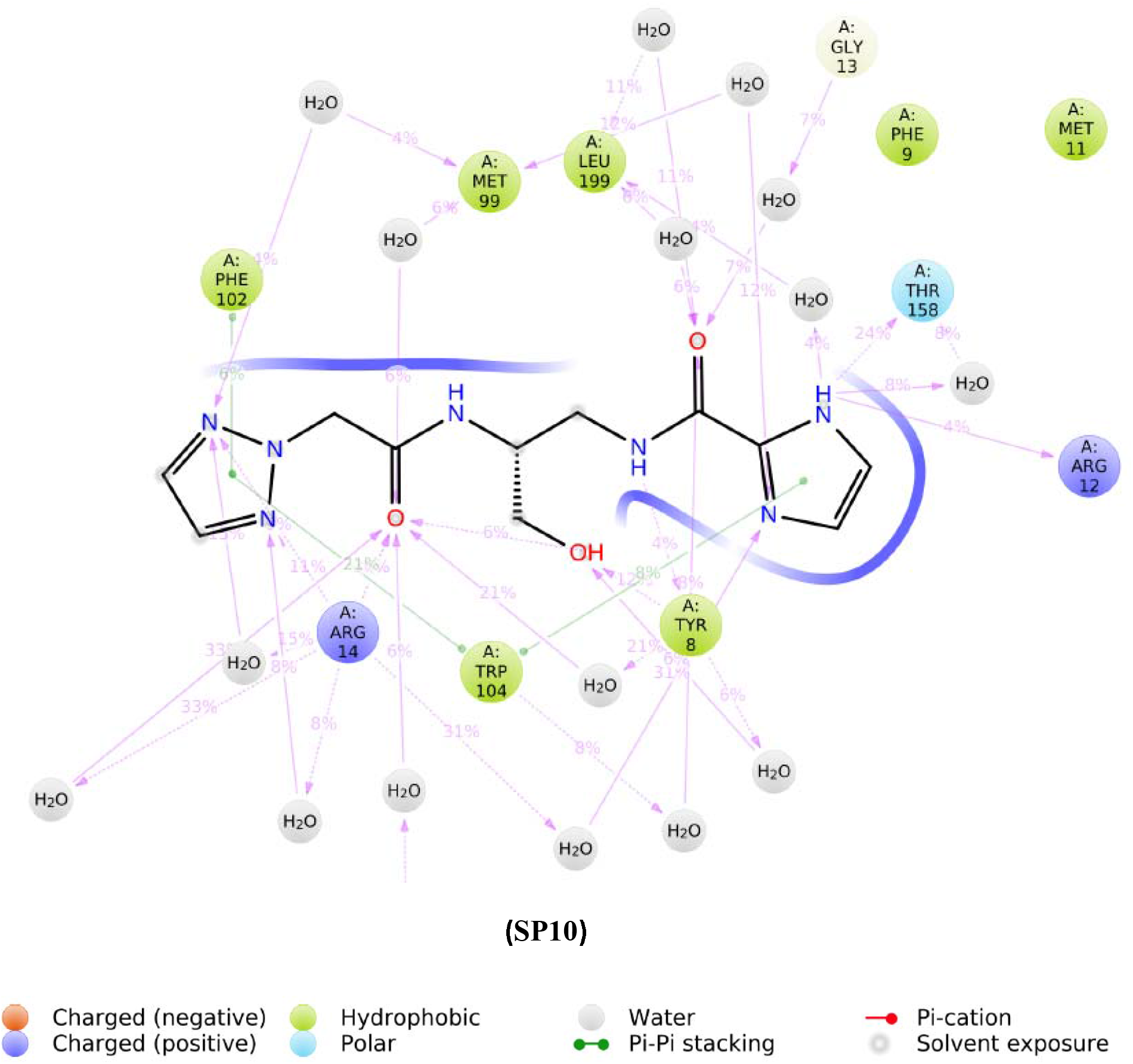
The protein-ligand contact diagram of **SP1, SP2 and SP10** ligands with human hematopoietic prostaglandin D2 synthase enzyme.

## Conclusion

This work helps in identifying a more effective drug candidate against the human hPGDS by performing pharmacophore-based virtual screening as a measuring tool. The pharmacophore-based virtual screening, molecular docking, MM_GBSA, ADME property analysis combinedly concluded with four ligands (**SP1**, **SP2**, **SP5**, and **SP10**) which have good docking score and ligand-receptor interaction in comparison to the compound reported in the literature and available in the market like **TFC-007**, **HPGDS inhibitor I**, **HQL-79** and **TAS-204**. But **2CVD-SP5** complex was not showing good stability in the ligand-receptor RMSD study of MD simulation. The outcome of this study concludes with three ligands (**SP1**, **SP2**, and **SP10**) which shows a good range of H-bonding due to H-bond donor and acceptor groups, pi-pi stacking due to the presence of ring aromatic compound, pi-cation, and multiple solvent exposures. The MD simulations validated our assumptions that **SP1**, **SP2**, and **SP10** ligands have better interactions and strong binding affinity with the human hPGD2 enzyme. Further, in vitro analysis followed by its in vivo testing may help in proving **SP1**, **SP2**, and **SP10** ligands as a better inhibitor of hPGD2.

## Supplementary Material

Supplementary material associated with this article can be found in the online version.

## Notes

The authors declare no competing financial interest.

## Acknowledgments

Mr. Satyajit is grateful to MHRD, India for financial support. The authors also acknowledge Mr. Vinod Deveraji (Application Scientist from Schrodinger, Banglore) for their technical assistance.

