## Supplemental data for "Identification of Potential inhibitors for Hematopoietic Prostaglandin D2 Synthase: Computational Modeling and Molecular Dynamics Simulations"

**Abstract**

To design a new therapeutic agent for Hematopoietic Prostaglandin D_2_ synthase (hPGDS), a set of 60 molecules with different molecular scaffolds were (range of pIC_50_ values are from 8.301 to 3.932) considered to create a pharmacophore model. Further, identification of potential hPGDS inhibitors were carried out by using virtual screening with different databases (from 15,74,182 molecules). The Molecular screening was performed using different sequential methods right from Pharmacophore based virtual screening, **molecular docking, MM-GBSAstudies, ADME property analysis and molecular dynamics simulations using** Maestro11.9 software**. Based on the best pharmacophore model (**ADRR_1), the resultant set of 18,492 molecules were screened. The preliminarily screened molecules were subjected to molecular docking (PDB_ID: 2CVD) methods**. A set of 27 molecules was screened from the resultant molecular docking outcomes (360 molecules) based on** binding free energy (ΔG_bind_) and Lipinski’s rule of five. Out of 27 molecules, 4 were selected visual data analysis and further subjected to molecular dynamics (MD) simulation study. Outcomes of the present study conclude with three new proposed molecules (**SP1, SP2** and **SP10)** which show a good range of interaction with human hPGDS enzyme in comparison to the marketed compounds i.e., **HQL-79**, **TFC-007**, **HPGDS inhibitor I** and **TAS-204**.

### To whom correspondence should be addressed:

*

[# Current address]: Department of Biotechnology, Indian Institute of Technology, Kharagpur-721302, India.

**Supporting Information**

Table S1 (A): Structures and biological activities of indole^a^ based Potential Hematopoietic Prostaglandin D2 Synthase inhibitor.

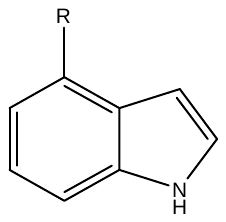

(A)

| Code | R | IC50 | Activity* |
| --- | --- | --- | --- |
| A1 | 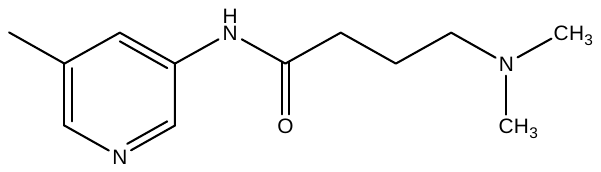 | 0.180 | 6.745 |
| A2 | 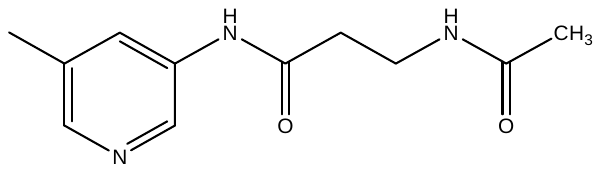 | 0.300 | 6.523 |
| A3 | 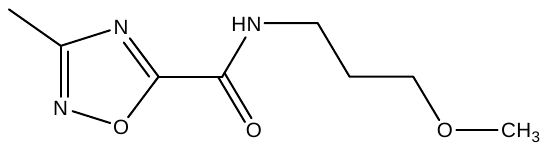 | 0.170 | 6.770 |
| A4 | 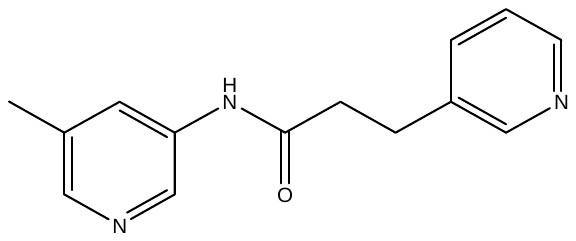 | 0.026 | 7.585 |
| A5 | 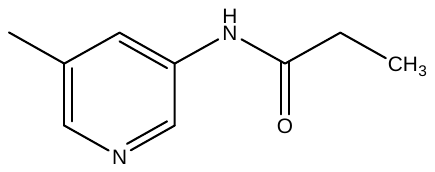 | 0.032 | 7.495 |
| A6 | 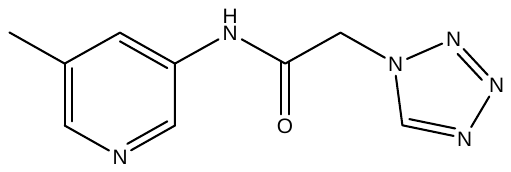 | 0.043 | 7.367 |
| A7 | 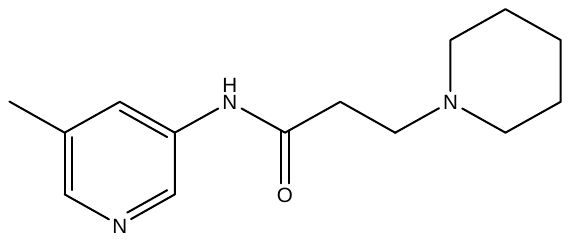 | 0.130 | 6.886 |
| A8 | 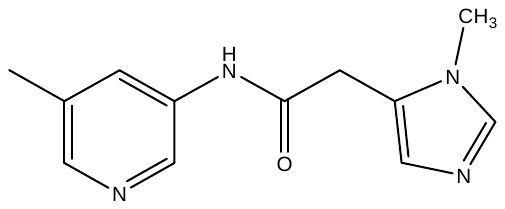 | 1.200 | 5.921 |

^a^[Edfeldt, 2015]

*Calculated by taking log_10_ of IC_50_ value as discussed in Data set preparation section

Table S1 (B): Structures and biological activities of pyridine ^a^ based Potential Hematopoietic Prostaglandin D2 Synthase inhibitor.

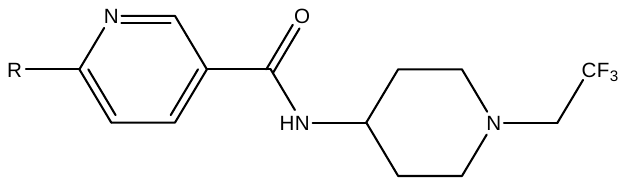

(B)

| Code | R | IC50 | Activity* |
| --- | --- | --- | --- |
| B9 | 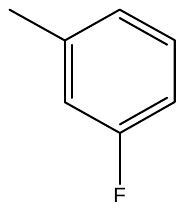 | 0.200 | 6.699 |
| B10 | 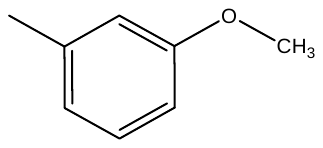 | 0.060 | 7.222 |
| B11 | 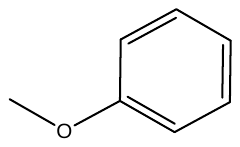 | 0.031 | 7.509 |
| B12 | 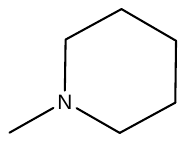 | 0.043 | 7.367 |
| B13 | 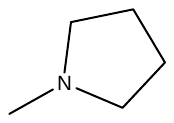 | 0.230 | 6.638 |
| B14 | 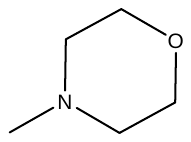 | 0.320 | 6.495 |
| B15 | 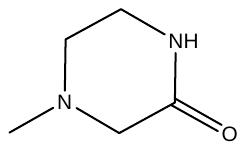 | 0.590 | 6.229 |
| B16 | 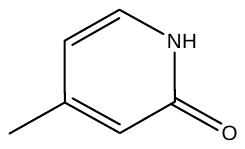 | 0.070 | 7.155 |
| B17 | 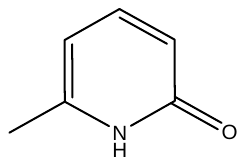 | 0.071 | 7.149 |

^a^[Edfeldt, 2015].

*Calculated by taking log_10_ of IC_50_ value as discussed in Data set preparation section

Table S1 (C): Structures and biological activities of benzaldehyde^b^ based Potential Hematopoietic Prostaglandin D2 Synthase inhibitor.

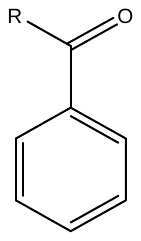

(C)

| Code | R | IC50 | Activity* |
| --- | --- | --- | --- |
| C18 | 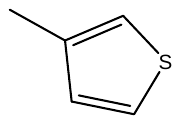 | 11.400 | 4.943 |
| C19 | 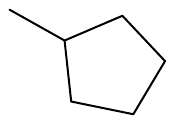 | 16.000 | 4.796 |
| C20 | 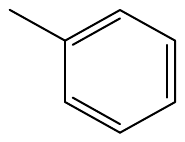 | 62.200 | 4.206 |
| C21 | 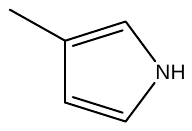 | 87.500 | 4.058 |
| C22 | 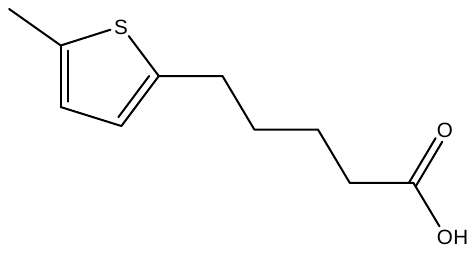 | 117.000 | 3.932 |

^b^[Weber, 2010].

*Calculated by taking log_10_ of IC_50_ value as discussed in Data set preparation section

Table S1 (D): Structures and biological activities of thiophene^c^ based Potential Hematopoietic Prostaglandin D2 Synthase inhibitor.

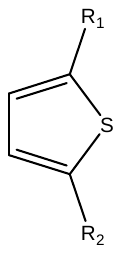

(D)

| Code | R1 | R1 | IC50 | Activity* |
| --- | --- | --- | --- | --- |
| D23 | 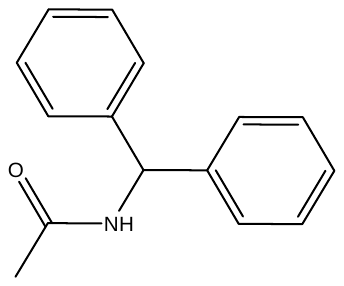 | 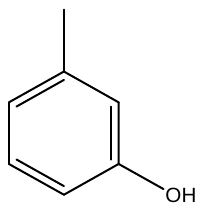 | 0.700 | 6.155 |
| D24 | 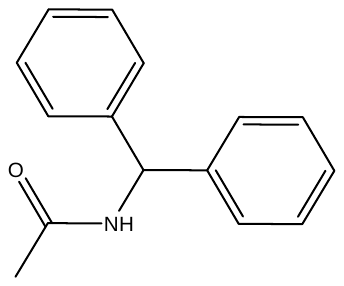 | 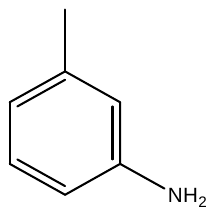 | 1.900 | 5.721 |
| D25 | 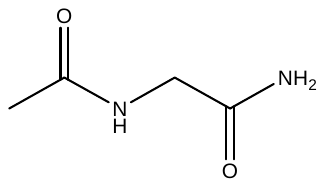 | 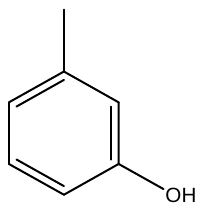 | 8.000 | 5.097 |
| D26 | 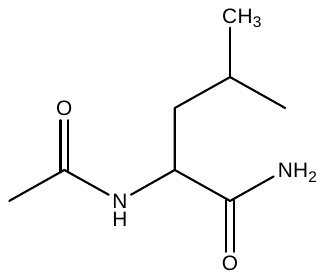 | 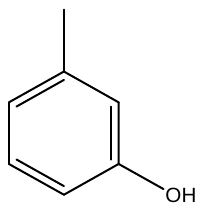 | 7.100 | 5.149 |
| D27 | 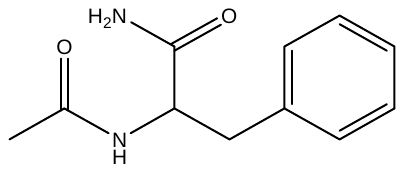 | 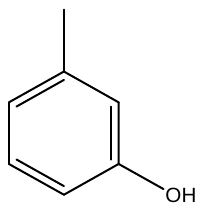 | 3.700 | 5.432 |
| D28 | 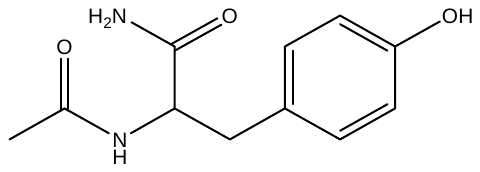 | 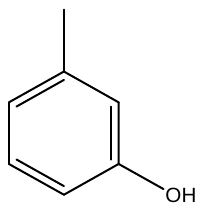 | 1.300 | 5.886 |
| D29 | 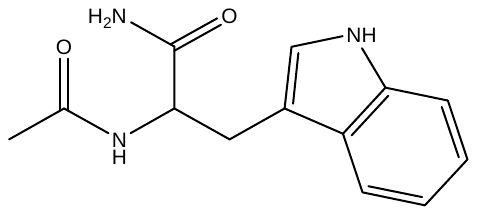 | 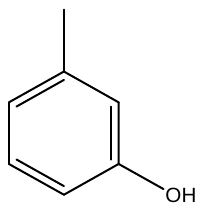 | 2.100 | 5.678 |

^c^[Christ, 2010]

*Calculated by taking log_10_ of IC_50_ value as discussed in Data set preparation section

Table S1 (E): Structures and biological activities of benzimidazole^d^ based Potential Hematopoietic Prostaglandin D2 Synthase inhibitor.

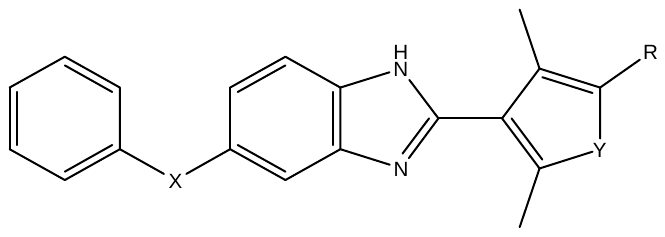

(E)

| Code | R | X | Y | IC50 | Activity* |
| --- | --- | --- | --- | --- | --- |
| E30 | CN | -C(O)- | NH | 0.540 | 6.268 |
| E31 | 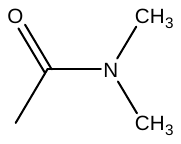 | -C(O)- | NH | 0.256 | 6.592 |
| E32 | 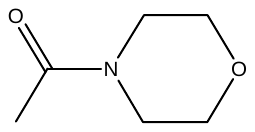 | -C(O)- | NH | 0.590 | 6.229 |
| E33 | 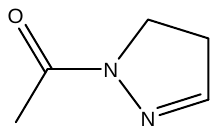 | -C(O)- | NH | 0.063 | 7.201 |
| E34 | 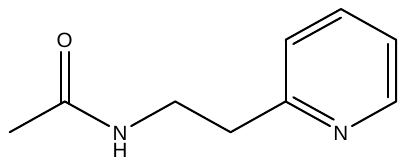 | -C(O)- | NH | 0.072 | 7.143 |
| E35 | 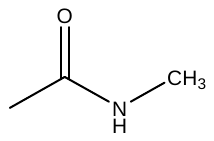 | -C(O)- | NH | 0.077 | 7.114 |
| E36 | COOH | -C(O)- | NH | 0.436 | 6.361 |
| E37 | CONH_2_ | -C(O)- | NH | 0.100 | 7.000 |
| E38 | 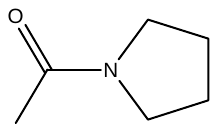 | -C(O)- | NH | 0.351 | 6.455 |
| E39 | 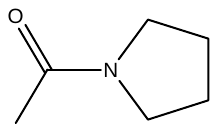 | -O- | NH | 0.614 | 6.212 |
| E40 | 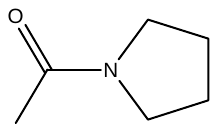 | -C(O)- | O | 0.178 | 6.750 |
| E41 | CONH_2_ | -O- | O | 0.109 | 6.963 |

^d^[Thurairatnam, 2012].

*Calculated by taking log_10_ of IC_50_ value as discussed in Data set preparation section

Table S1 (D): Structures and biological activities of pyrimidine ^d^ based Potential Hematopoietic Prostaglandin D2 Synthase inhibitor.

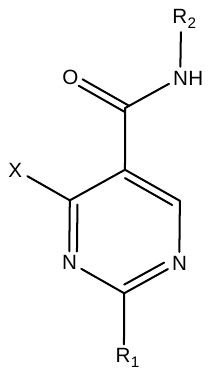

(F)

| Code | X | R1 | R2 | IC50 | Activity* |
| --- | --- | --- | --- | --- | --- |
| F42 | H | 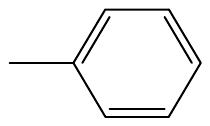 | 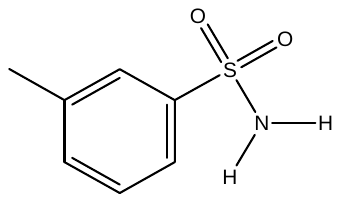 | 0.005 | 8.301 |
| F43 | H | 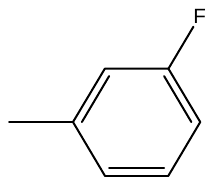 | 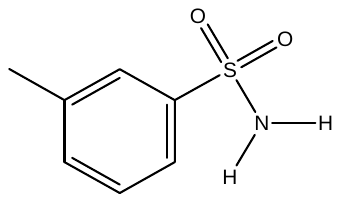 | 0.031 | 7.509 |
| F44 | H | 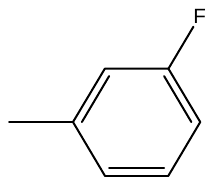 | 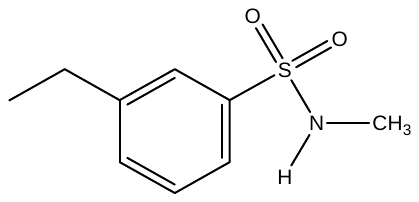 | 0.009 | 8.022 |
| F45 | H | 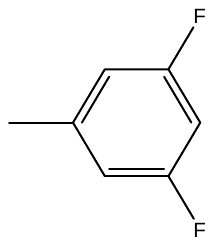 | 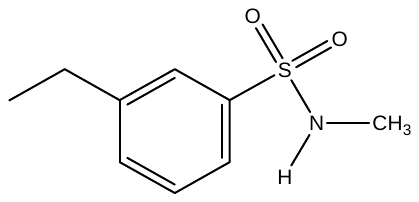 | 0.049 | 7.310 |
| F46 | CH_3_ | 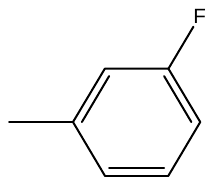 | 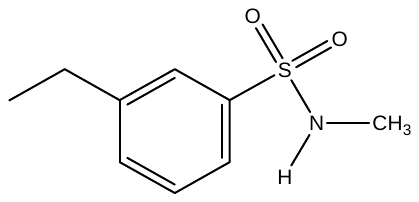 | 0.106 | 6.975 |
| F47 | H | 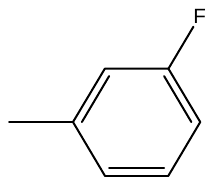 | 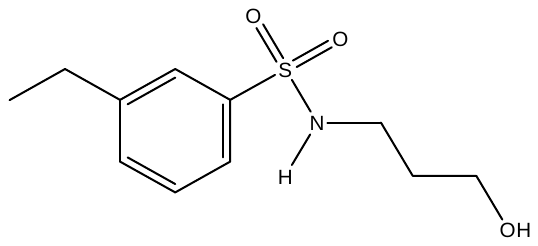 | 0.022 | 7.658 |
| F48 | CH_3_ | 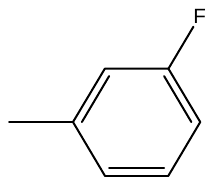 | 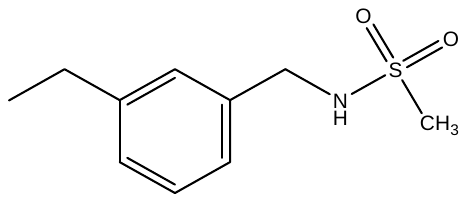 | 0.049 | 7.310 |
| F49 | H | 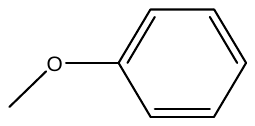 | 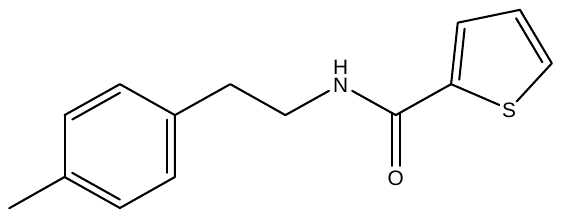 | 0.141 | 6.851 |
| F50 | H | 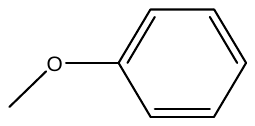 | 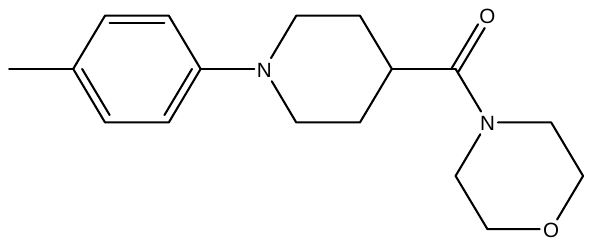 | 0.280 | 6.553 |
| F51 | H | 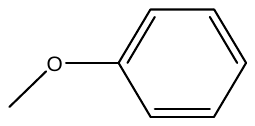 | 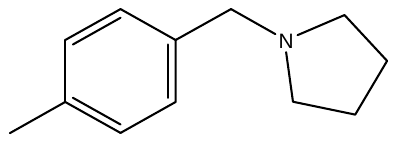 | 0.728 | 6.138 |
| F52 | H | 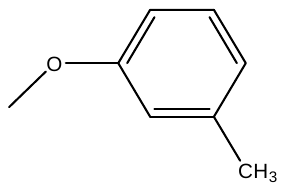 | 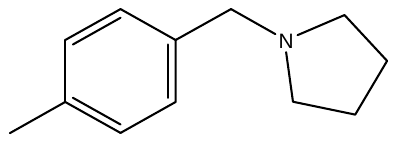 | 2.360 | 5.627 |
| F53 | H | 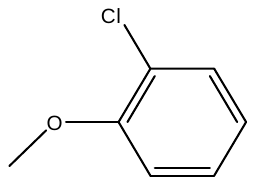 | 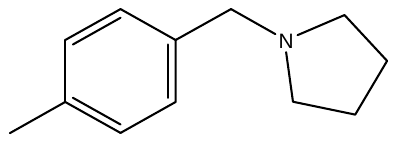 | 2.990 | 5.524 |
| F54 | CH_3_ | 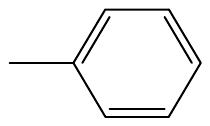 | 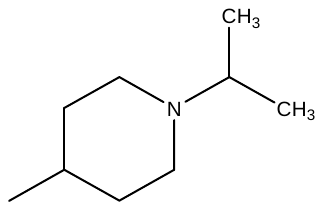 | 0.162 | 6.790 |
| F55 | H | 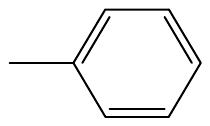 | 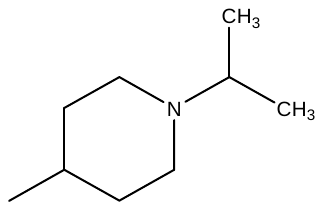 | 0.006 | 8.198 |
| F56 | NH_2_ | 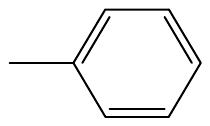 | 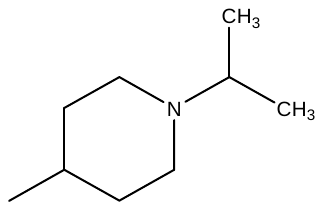 | 0.219 | 6.660 |
| F57 | OH | 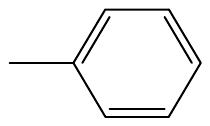 | 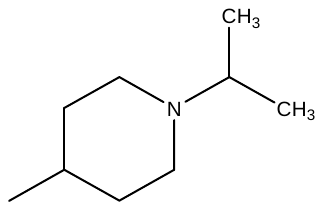 | 1.000 | 6.000 |
| F58 | OH | 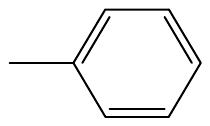 | 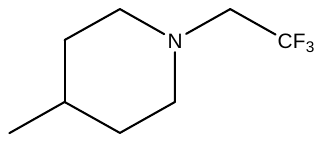 | 1.000 | 6.000 |
| F59 | H | 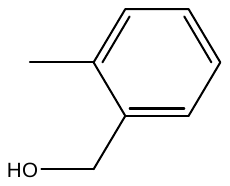 | 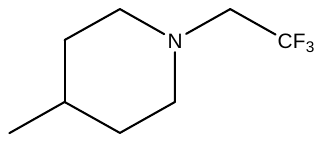 | 1.010 | 5.996 |
| F60 | H | 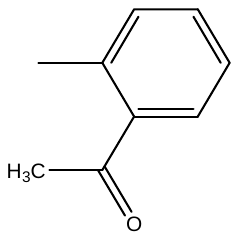 | 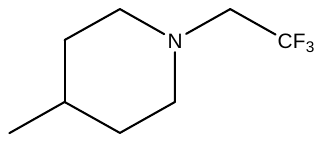 | 9.460 | 5.024 |

^d^[Thurairatnam, 2012].

*Calculated by taking log_10_ of IC_50_ value as discussed in Data set preparation section

**Table S2:** Structures of Hematopoietic Prostaglandin D2 Synthase inhibitors (Molecules screened by ADME Screening).

| **Code** | **Structure** |
| --- | --- |
| **SP1** | 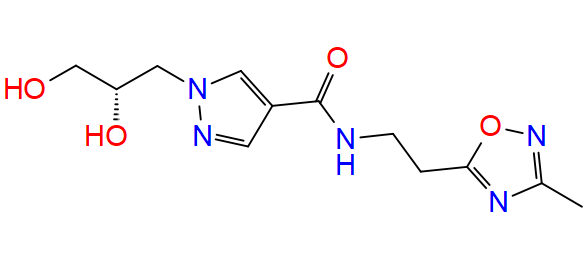 |
| **SP2** | 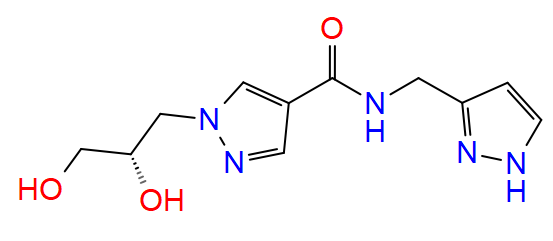 |
| **SP5** | 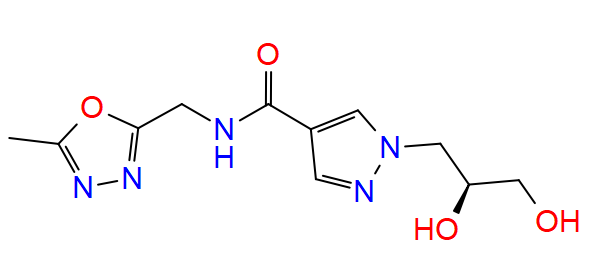 |
| SP7 | 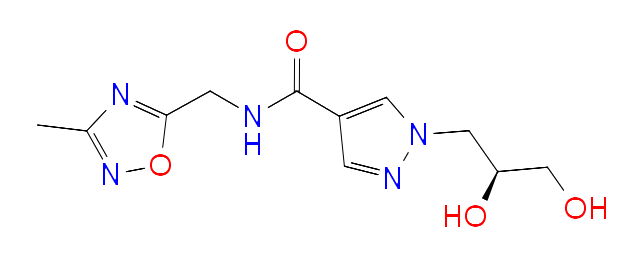 |
| SP9 | 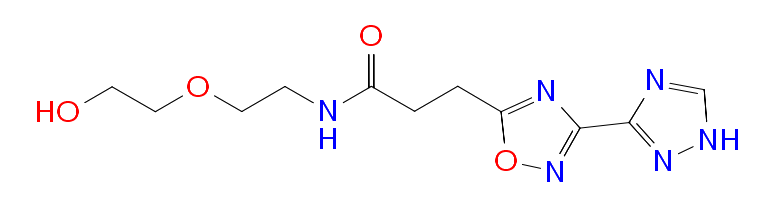 |
| **SP10** | 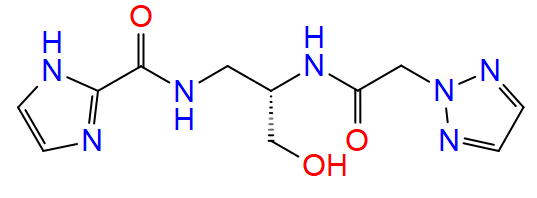 |
| SP11 | 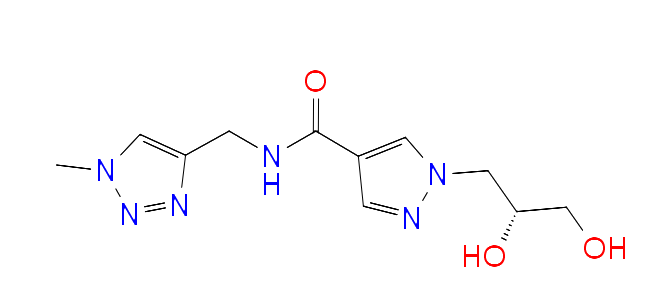 |
| SP12 | 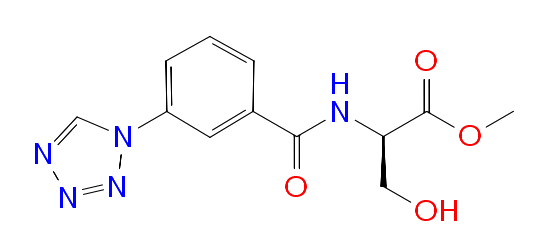 |
| SP13 | 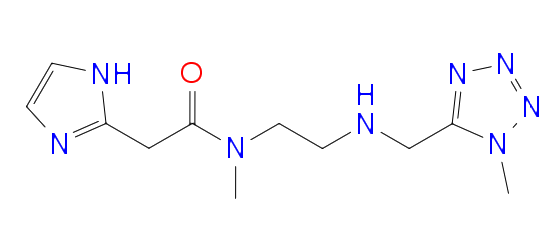 |
| SP14 | 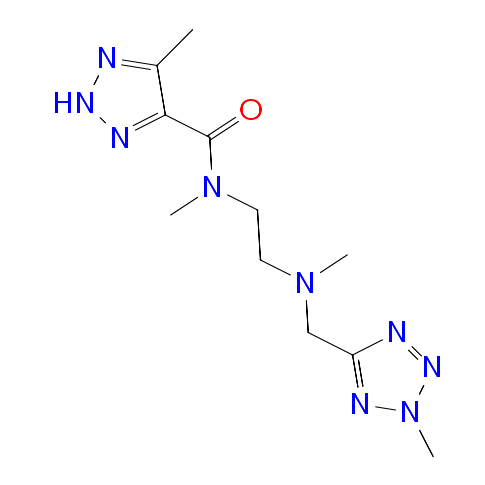 |
| SP15 | 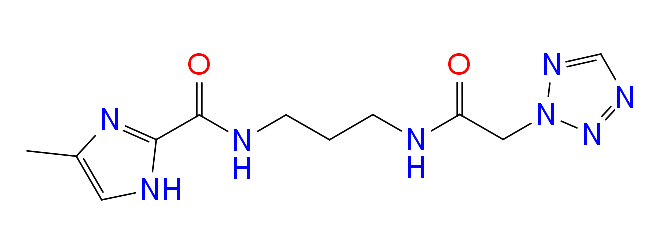 |
| SP17 | 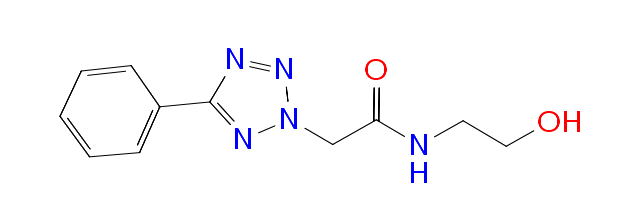 |
| SP18 | 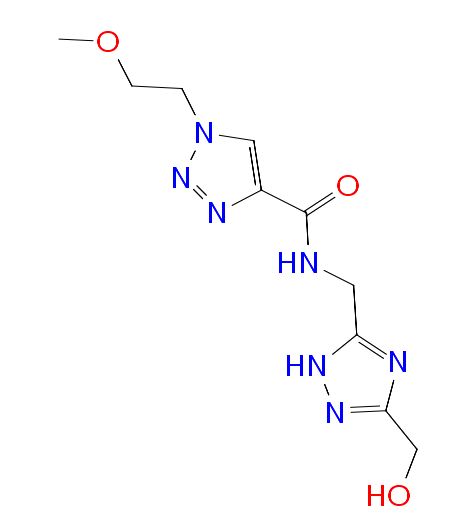 |
| SP19 | 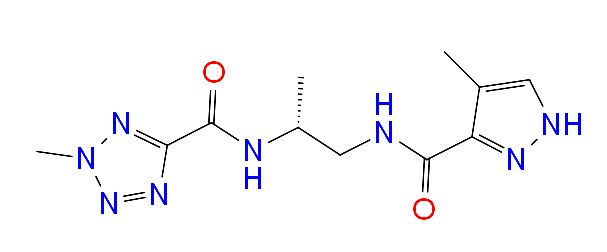 |
| SP20 | 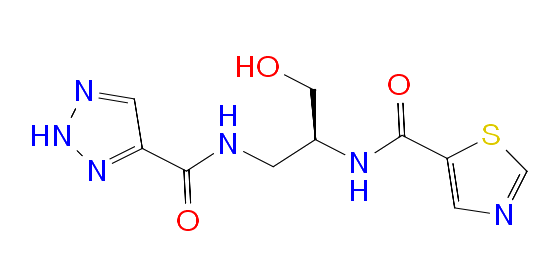 |
| SP21 | 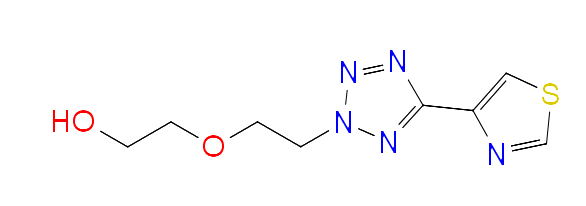 |
| SP22 | 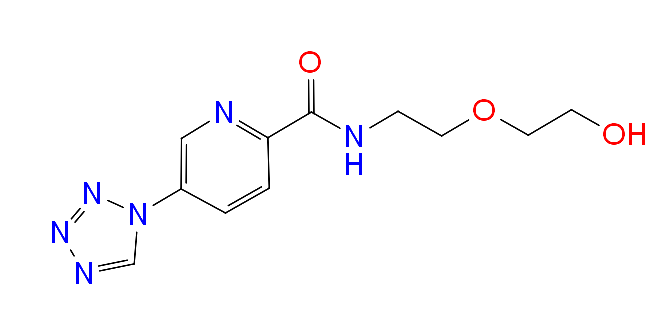 |
| SP23 | 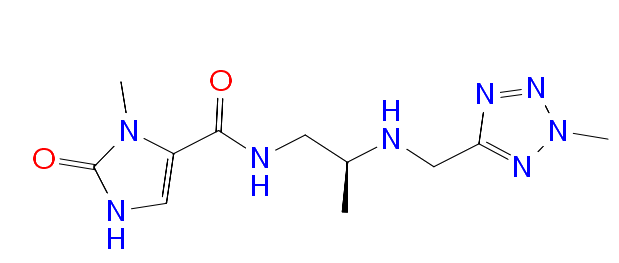 |
| SP24 | 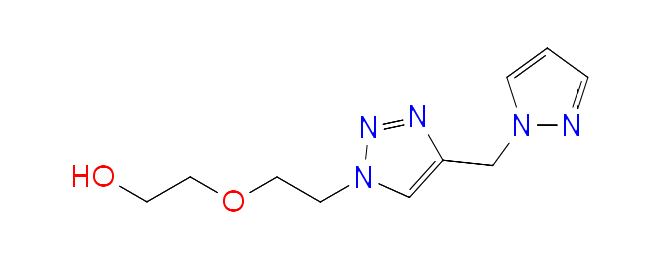 |
| SP26 | 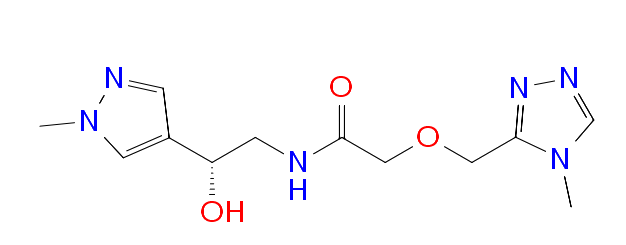 |
| SP27 | 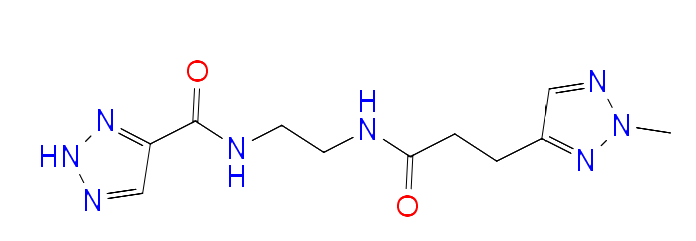 |
| SP29 | 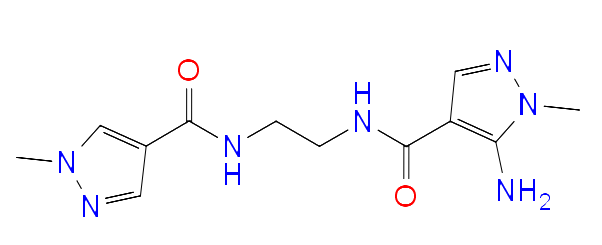 |
| SP32 | 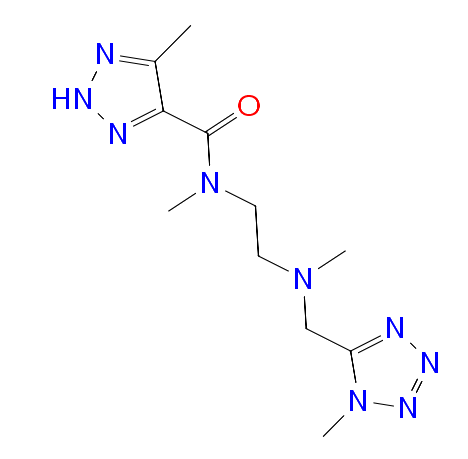 |
| SP33 | 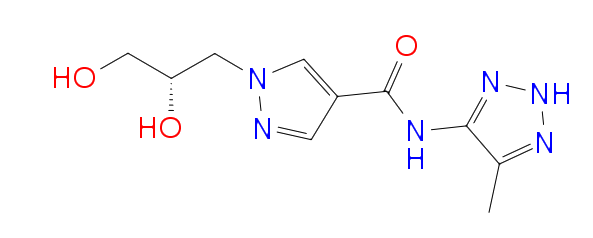 |
| SP35 | 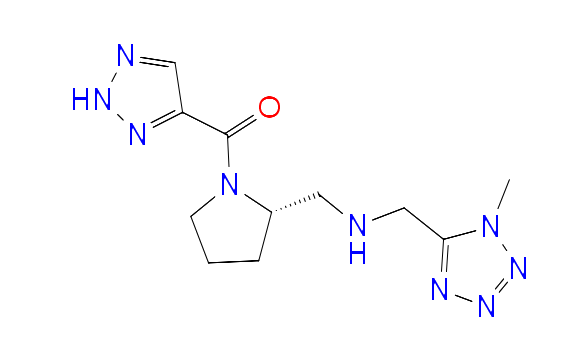 |
| SP37 | 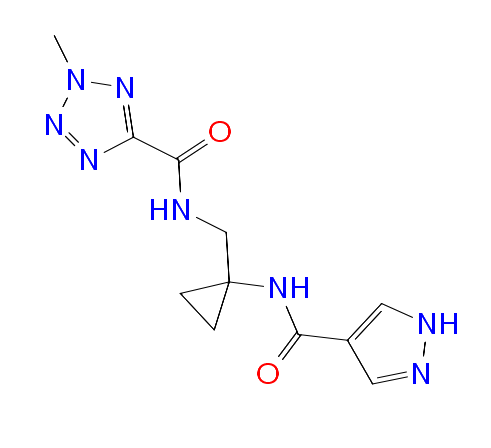 |
| SP38 | 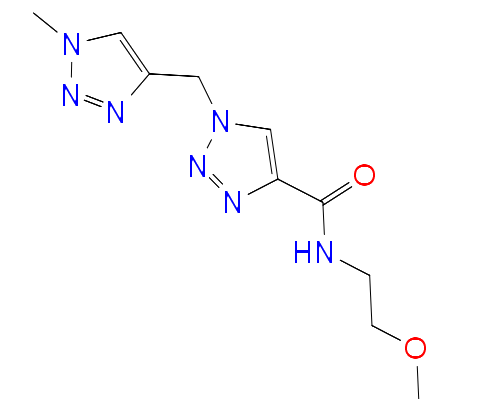 |

**Fig.S1:** Residues interact with the ligand in each trajectory frame.

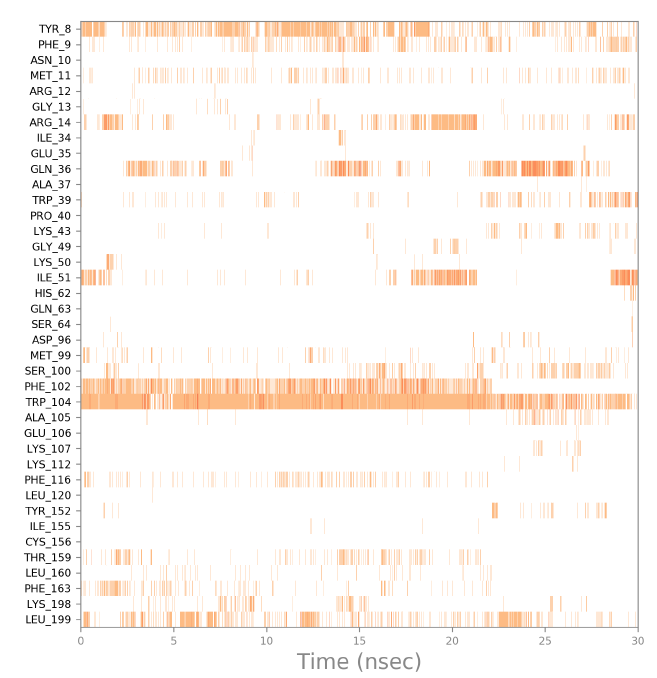

**(SP1)**

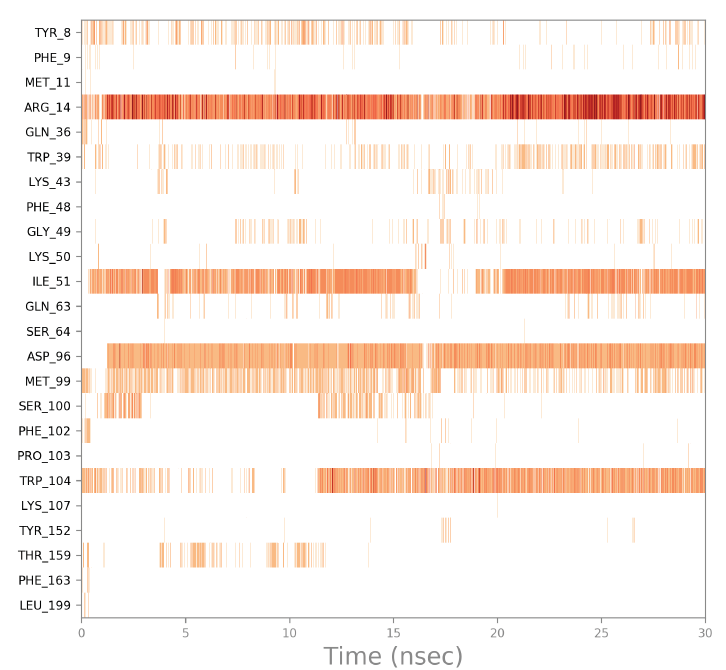

**(SP2)**

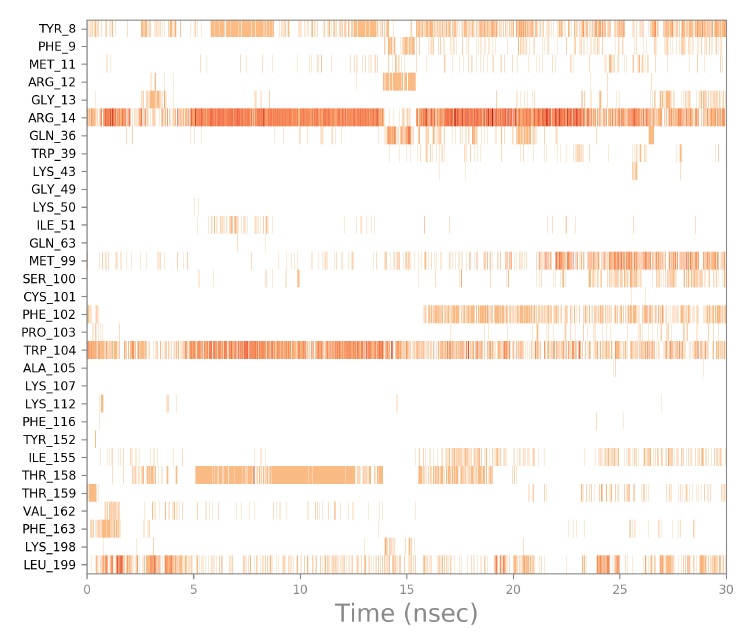

**(SP10)**
